# Composite Representations Lead to Multiple Geometrical Approximations to the Structure of Musical Pitch

**DOI:** 10.1101/2023.06.13.544763

**Authors:** Raja Marjieh, Thomas L. Griffiths, Nori Jacoby

**Affiliations:** Department of Psychology, Princeton University, USA; Department of Computer Science, Princeton University, USA; Max Planck Institute for Empirical Aesthetics, Germany; Department of Psychology, Cornell University, USA

**Author notes:** Corresponding author: Raja Marjieh. Data and code: https://osf.io/nx586/?view_only=a0f74c9269a4448f821aadccf76ba175. Equal Contribution (Thomas L. Griffiths, Nori Jacoby).

**Keywords:** pitch perception, representations, octave equivalence, singing

## Abstract

Pitch perception is central for both speech and music, and the representation of musical pitch has intrigued scholars for centuries. In his seminal work, Roger Shepard proposed a series of increasingly complex geometrical models to approximate the psychological representation of pitch, most famously the pitch helix which is often used in textbooks.

The pitch helix represents the logarithmic scaling of the periodicity of tones and the similarity between tones separated by an octave. However, support for geometrical models of musical pitch derives from studies that are small-scale and sometimes contradictory.

Moreover, research suggests that the representation of pitch is influenced by task context and expertise raising the question of whether any single integrated geometric approximation is sufficient. Using multi-dimensional scaling analysis, we revisit this problem through nine experiments involving participants with varied levels of musical expertise (*N* = 592) and paradigms covering both perception and production, as well as implicit and explicit measures of perceptual similarity. We show that, depending on task and musical experience, the best geometrical approximation can exhibit an array of structures ranging from linear to double-helical structures, providing strong evidence that a simple helical model, or in fact any fixed geometrical model, cannot explain the data. To explain these large variations, we then show that they can be captured by different reweightings of a small set of perceptual factors suggesting a composite representation.

These findings highlight the importance of examining diverse tasks and populations to address the classic question of how perceptual representations are organized.

---

Pitch is one of the most extensively studied psychological phenomena due to its crucial role in both music and speech perception (Attneave & Olson, 1971; Koelsch & Siebel, 2005; Peretz & Zatorre, 2005; Levitin, 2006; Plack et al., 2006; Zatorre et al., 2007; McDermott & Oxenham, 2008; Patel, 2010; Wong et al., 2012; McPherson & McDermott, 2018; Oxenham, 2018; Jacoby et al., 2019). Research on pitch typically focuses on two complementary questions. The first major question concerns the computational and neural processes underlying the local transformation from a given sound sensation to pitch perception (Plack et al., 2006; McDermott & Oxenham, 2008; Oxenham, 2018). In this area, researchers have explored the integration of various cues (Von Békésy, 1960; Rose et al., 1967; Patterson, 1976; Shamma, 1985; Cedolin & Delgutte, 2010; Bentsen et al., 2011), including resolved and unresolved harmonics (Plomp, 1967; Shackleton & Carlyon, 1994; Lau et al., 2017). Moreover, empirical studies have investigated the neural correlates of pitch processing (Schulze et al., 2002; Bendor & Wang, 2005; Schnupp et al., 2011; Norman-Haignere et al., 2013; Feng & Wang, 2017), while computational modeling has provided insights into the biological mechanisms involved in this transformation (de Cheveigné, 2010).

The second major question concerns how individual pitch percepts are combined to form larger perceptual structures, such as pitch contours in speech perception, which convey prosodic information, or lexical distinctions in tone languages. In music, individual pitch percepts are organized into scales that reference tonal centers (Krumhansl & Kessler, 1982; Fogel et al., 2015), ultimately creating melodies (Cuddy et al., 1981; Dowling, 1990; Temperley, 2008; Anglada-Tort et al., 2023) and complete musical compositions (Zatorre et al., 1994; Cheung et al., 2019; Zatorre, 2024). Key to this question are the underlying global representations that organize the relationships between different pitch percepts and the way they can be effectively modeled (Shepard, 1982a, 1982b; Patel, 2003).

Perhaps the most well-known of these structures is the pitch helix (Figure 1A-B; see also Supplementary Figure A12). This model, proposed by Shepard (Shepard, 1978, 1982a, 1982b; though see also Ruckmick, 1929) accounts for two key features of pitch in Western music. First, pitch is organized uniformly on a logarithmic scale (i.e., a log-linear line; Jacoby et al., 2019). Tones with slower periodicity are perceived as “low”, while those with faster periodicity are perceived as “high” (ANSI, 1960). Additionally, Western music uses octave equivalence, where tones with periodicities differing by an octave (a 2:1 ratio) share the same note name, or chroma, and are treated as musically similar (Shepard, 1964; Deutsch, 1973; Kallman, 1982; Shepard, 1982a; Demany & Armand, 1984; Hoeschele et al., 2012, 2013; Jacoby et al., 2019). The combination of these features—pitch height and octave equivalence—led to the hypothesis that musical pitch representations could be approximated by a helix (Shepard, 1982a). Since then, the pitch helix and its sub-components have become a standard topic in many perception and music textbooks (e.g., Krumhansl, 2001; Plack et al., 2006; Tymoczko, 2006; Wolfe et al., 2006; Goldstein, 2008; Plack, 2018), as well as modern research and review articles (e.g., Warren et al., 2003; Trainor and Unrau, 2011; Hoeschele et al., 2012; Milne and Holland, 2016; Jacoby et al., 2019; Marjieh, Sucholutsky, et al., 2024).

**Figure 1.**
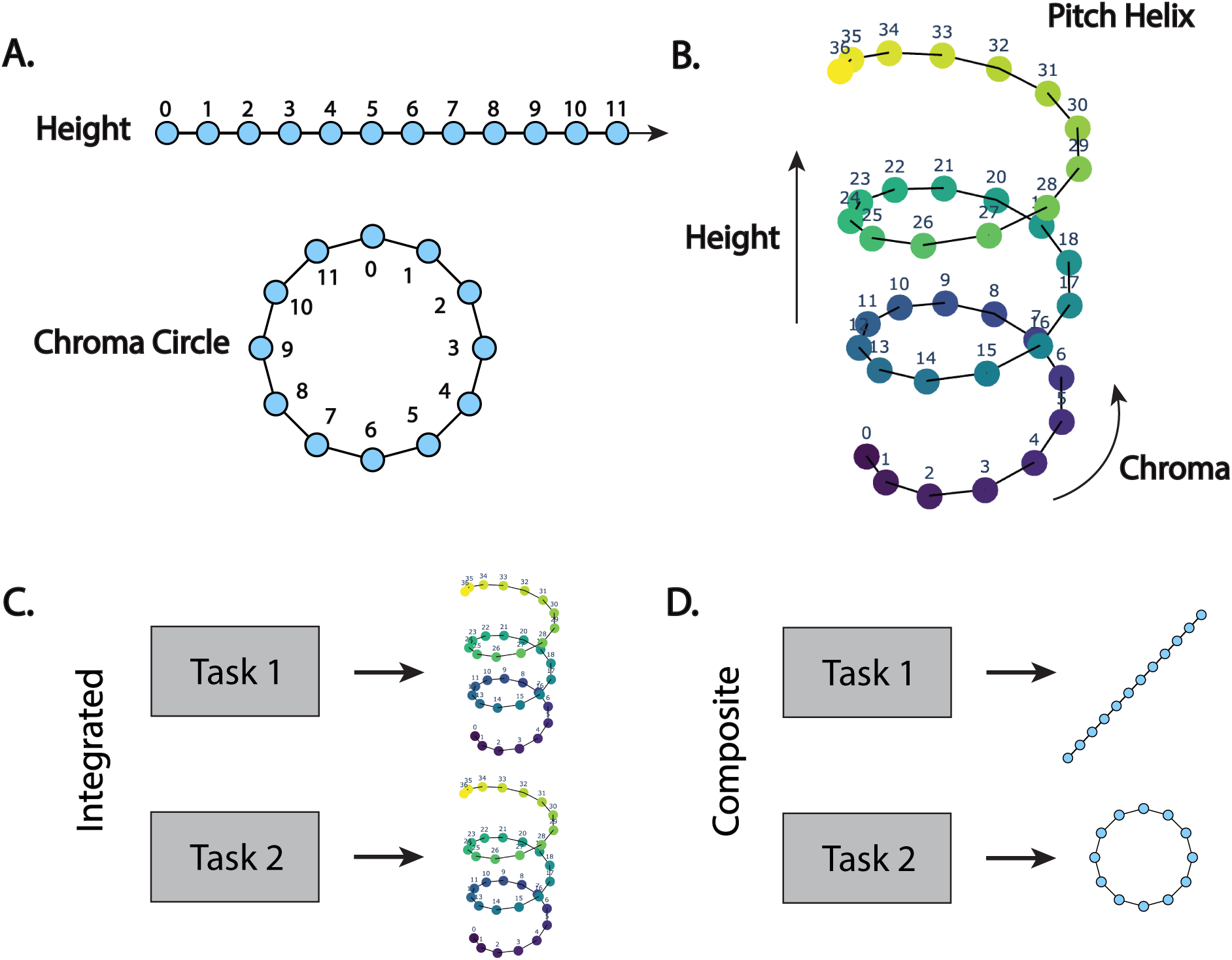
A geometrical model of the representation of musical pitch. **A.** Pitch height and pitch chroma components. **B.** The combined pitch helix. Numbers indicate semitonal pitch values relative to a generic base labeled as 0. **C.** An integrated representation: the pitch helix is detectable across tasks. **D.** A composite representation: different components are activated to different degrees across tasks.

Despite its success, there are reasons to question whether the pitch helix is an adequate model of musical pitch. First, octave equivalence is still actively debated in the literature, both with respect to its dependence on biological mechanisms and its prevalence in the population (Deutsch, 1972; Warren et al., 2003; Briley et al., 2013; Hoeschele et al., 2013; Jacoby et al., 2019; Pressnitzer and Demany, 2019; Regev et al., 2019). Second, the pitch helix fails to capture other significant musical intervals, in particular, the perfect fifth and its inversion, the perfect fourth, which constitute the circle of fifths. To accommodate these, Shepard (1982b) proposed a series of increasingly complex, though perhaps less well-known, geometrical models based on a double-helix^1^ structure wrapped around other structures such as a line or a torus (see Supplementary Figure A12). Alternative structures have also been proposed to capture the influence of musical context on pitch relations, notably by Krumhansl (1979) (see also Krumhansl and Kessler (1982) for more abstract models of musical key relations). However, empirical evidence for these models suffers from multiple methodological limitations, including small sample sizes, reliance on highly trained musicians, restriction to a single task, the use of unsuitable paradigms, and the dependence on fixed priming schemes (see next section for additional details).

From a theoretical standpoint, a geometrical model such as the pitch helix also implies an *integrated* (or unified) representation that serves as a substrate for cognitive processes across tasks (e.g., pitch imitation or melody identification, Jacoby et al., 2019; Deutsch, 1972). Integrated representations have proven highly productive in understanding other perceptual spaces, such as color, where they have also led to significant applications (e.g. CIELAB; Schanda, 2007). If this is indeed the case, then one would expect to detect such a structure by examining behavioral data from any sufficiently expressive and powered task (Figure 1C). This perspective contrasts an alternative view, namely, that the representation of pitch is *composite* and reflects multiple distinct factors (e.g., logarithmic frequency scaling and octave equivalence) that are activated to different degrees depending on task demands (Figure 1D). While in some tasks the congruence of such factors may globally align with a geometrical model such as the pitch helix, this may not always be the case. Although subtle, this difference has important empirical and theoretical implications. From an empirical perspective, this necessitates an evaluation of geometrical models of pitch across multiple tasks in parallel which is rarely done as noted earlier. From a modeling perspective, this implies that a more adequate theoretical formulation would explicitly incorporate configurable perceptual factors rather than directly relying on the global geometrical model that reflects a specific weighting of those factors. Such a perspective has some precedence in preliminary work by Shepard (1982b), and it also aligns with studies on local pitch perception (i.e., the local transformation from a given sound sensation to pitch perception) that deploy multiple tasks to isolate distinct mechanisms involved in the process (e.g., McPherson and McDermott, 2018).

In this work, we address this gap in the literature by jointly probing the representation of pitch across tasks and experience groups and evaluating how it explicitly translates into geometrical models. In addition, we address previous methodological limitations by applying new techniques from large-scale online experiments (Anglada-Tort et al., 2023; Marjieh, Jacoby, et al., 2024), enabling a fresh look at the global representation of musical pitch. Our approach, inspired by multi-dimensional scaling (MDS; Shepard, 1980) and representational similarity analysis (RSA; Kriegeskorte et al., 2008), covers both production-based and evaluative paradigms. Crucially, these paradigms allow us to elicit both implicit (singing responses) and explicit (direct ratings) measures of perceptual similarity. If the pitch helix, or any other structure, serves as a suitable geometrical model of pitch, then we would expect it to be detectable across the various tasks and experience groups. Alternatively, if this is not the case, then our goal is to provide a computational framework that can explain the observed variation in terms of a configurable set of perceptual factors.

The remainder of the paper proceeds as follows. First, we review empirical evidence on the global organization of musical pitch and its corresponding geometrical models.

Then, we introduce our behavioral paradigms and the general approach, followed by the methodological details. Next, we present our findings which reveal that the best geometrical approximations for the representation of pitch differ substantially across the different conditions, varying from a simple linear model to a complex double-helical one of the kind hypothesized by Shepard (1982a). These results challenge the idea of the pitch helix, as well as integrated geometrical models more broadly. To explain these variations, we then show how the majority of the variance in the human data can be explained by the reweighting of a small set of distinct perceptual factors which we identify. We conclude by discussing the implications of our findings for the psychology of pitch perception.

## Empirical Evidence for the Structure of Musical Pitch

### Pitch Height

Research has uncovered several findings that support the idea of pitch organization from low to high. The ear itself performs frequency decomposition, with high frequencies processed closer to the base of the cochlea (Von Békésy, 1960; Schnupp et al., 2011). This tonotopic organization is maintained in both the midbrain and primary auditory cortex (Winter et al., 2001; Patterson et al., 2002; Hall & Plack, 2009). Even non-experts are capable of making low-high judgments with considerable accuracy (Plack & Oxenham, 2005), and pitch perception follows a roughly logarithmic scale across a broad frequency range (Attneave & Olson, 1971; Moore, 1973; Jacoby et al., 2019). However, this mapping is not entirely straightforward, as pitch perception is influenced by multiple cues (Schnupp et al., 2011). Additionally, tonotopic regions including the auditory cortex do not respond robustly in the same way to tones of the same pitch with different timbres (Winter et al., 2001; Schulze et al., 2002; Hall & Plack, 2009) which indicates that they reflect lower-level, frequency-related aspects of tones rather than a fully invariant pitch representation (Nelken, 2011).

### Octave Equivalence

The findings on octave equivalence are more varied. Octave equivalence is fundamental to Western music theory (d’Arezzo, 1955; Drabkin, 2001; Aldwell et al., 2018), and octaves are common across songs from various cultures (Kuroyanagi et al., 2019; Mehr et al., 2019). However, the literature yields a complex picture with respect to its dependence on biological mechanisms, its prevalence among individuals with or without musical training, and its dependence on tasks (Humphreys, 1939; Shepard, 1964; Deutsch, 1973; Idson & Massaro, 1978; Kallman & Massaro, 1979; Kallman, 1982). Deutsch (1972) used a recognition paradigm to show that participants were able to reliably identify a famous tune when transposed into any of three given octave registers. However, when each note in the tune was independently sampled from a different octave register, recognition was severely diminished suggesting that octave equivalence cues were weaker than contour cues. Kallman (1982) used a similarity rating method testing three musicians, finding that only one of the three demonstrated a significant increase in perceived similarity for tones separated by an octave. Later, Demany and Armand (1984) used a melodic similarity paradigm, finding octave equivalence in children but noting a weaker effect in adults. More recently, Jacoby et al. (2019) used a singing paradigm, showing that octave equivalence is culturally contingent (absent among the Tsimane’, an indigenous Bolivian population) and varies within Western participants based on musical expertise. In contrast, Hoeschele and colleagues (Hoeschele et al., 2012, 2013) used an operant conditioning test with humans and chickadees, finding that only humans exhibited octave equivalence (see also Wagner et al., 2022, 2023, 2024). However, the degree to which octave equivalence is based on biological mechanisms remains highly debated (Pressnitzer & Demany, 2019), with consensus proving difficult due to the diverse range of paradigms, participant groups, and analytical methods used across studies.

Research on the neural basis of octave equivalence yields a nuanced picture as well. For instance, Warren et al. (2003) used fMRI to suggest distinct representations for chroma (anterior to the primary auditory cortex) and pitch height (posterior to the primary auditory cortex). In contrast, Briley et al. (2013), using EEG, found that chroma representation emerges only for complex tones, while pure tones primarily evoke representations of pitch height. More recently, Regev et al. (2019) reached a similar conclusion using EEG signal for deviance detection, showing that octave equivalence was not automatically detected with pure tones, even by expert musicians.

### The Pitch Helix and Other Geometrical Models

As for the the psychological research that directly addresses the question of the pitch helix and other geometrical models, it is surprising that it has largely relied on limited data and experimental designs. For instance, some studies that collected direct similarity judgments between tone pairs relied on small musician groups and yielded conflicting results (e.g., Kallman (1982) found evidence for octave equivalence in only one out of the three musicians tested, whereas Allen (1967) detected octave equivalence in data from ten musicians). Some relied exclusively on participants with significant musical experience (Krumhansl, 1979) which limits generalizability in light of evidence that auditory and pitch perception may depend on individual experience (Koelsch et al., 1999; Kishon-Rabin et al., 2001; Gaser & Schlaug, 2003; Fujioka et al., 2004; Peretz & Zatorre, 2005; Micheyl et al., 2006; Wong et al., 2007; Zatorre et al., 2007; Trainor & Corrigall, 2010; Dellacherie et al., 2011; François et al., 2013; Jacoby & Ahissar, 2013; Merzenich et al., 2013; Weiss et al., 2014; Liu et al., 2023). Additionally, studies using Shepard tones display octave equivalence by design as Shepard tones are physically identical for pitches separated by an octave, making it hard to draw inferences about the structure of the underlying helical representation (Shepard, 1978, 1982a). Notably, multiple studies that proposed alternative geometrical models confounded pitch perception with tonality. This is problematic as it is less surprising to see expertise differences in tasks involving tonality, a more complex musical feature, compared to pitch. For example, Shepard’s foundational work (Shepard, 1982a, 1982b) used probe tone ratings from Krumhansl and Shepard (1979) of how well a tone completes a scale and interpreted them as symmetric similarity scores between two pitch values. Similarly, Krumhansl’s seminal research on pitch similarity (Krumhansl, 1979) employed specific musical sequences before the comparison tones, like a diatonic C major scale, potentially priming participants to the initial sequence tones, as noted by Shepard (1982b). Later studies relied on a similar priming approach, and further conflated similarity with melodic consonance by asking participants to rate the “pleasantness” and “harmony” of the tone combinations (Amano, 1993).

In summary, many perception textbooks, handbooks, and articles (e.g., Krumhansl, 2001; Plack et al., 2006; Wolfe et al., 2006; Goldstein, 2008; Trainor and Unrau, 2011; Plack, 2018), use the pitch helix for describing aspects of pitch perception. However, evidence from multiple disparate studies suggests that pitch representations may be influenced by musical experience and task demands, which imply a more nuanced representation. In addition, previous psychological research that directly evaluates the pitch helix and other geometrical models suffers from various methodological limitations, including studying expertise and task separately, small sample sizes, and unsuitable paradigms that conflate factors such as tonality and consonance. These issues have led to a fragmented picture of pitch representation, making it difficult to determine whether pitch is represented as an integrated, task-general structure or as a composite of multiple distinct dimensions whose weight is task-dependent.

### A Holistic Test Across Tasks and Experience

To cover the different ways in which people process musical pitch, we considered three prominent behavioral paradigms (Figures 2A-C) that provide complementary forms of task engagement and applied them to both musician and non-musician participants.

**Figure 2.**
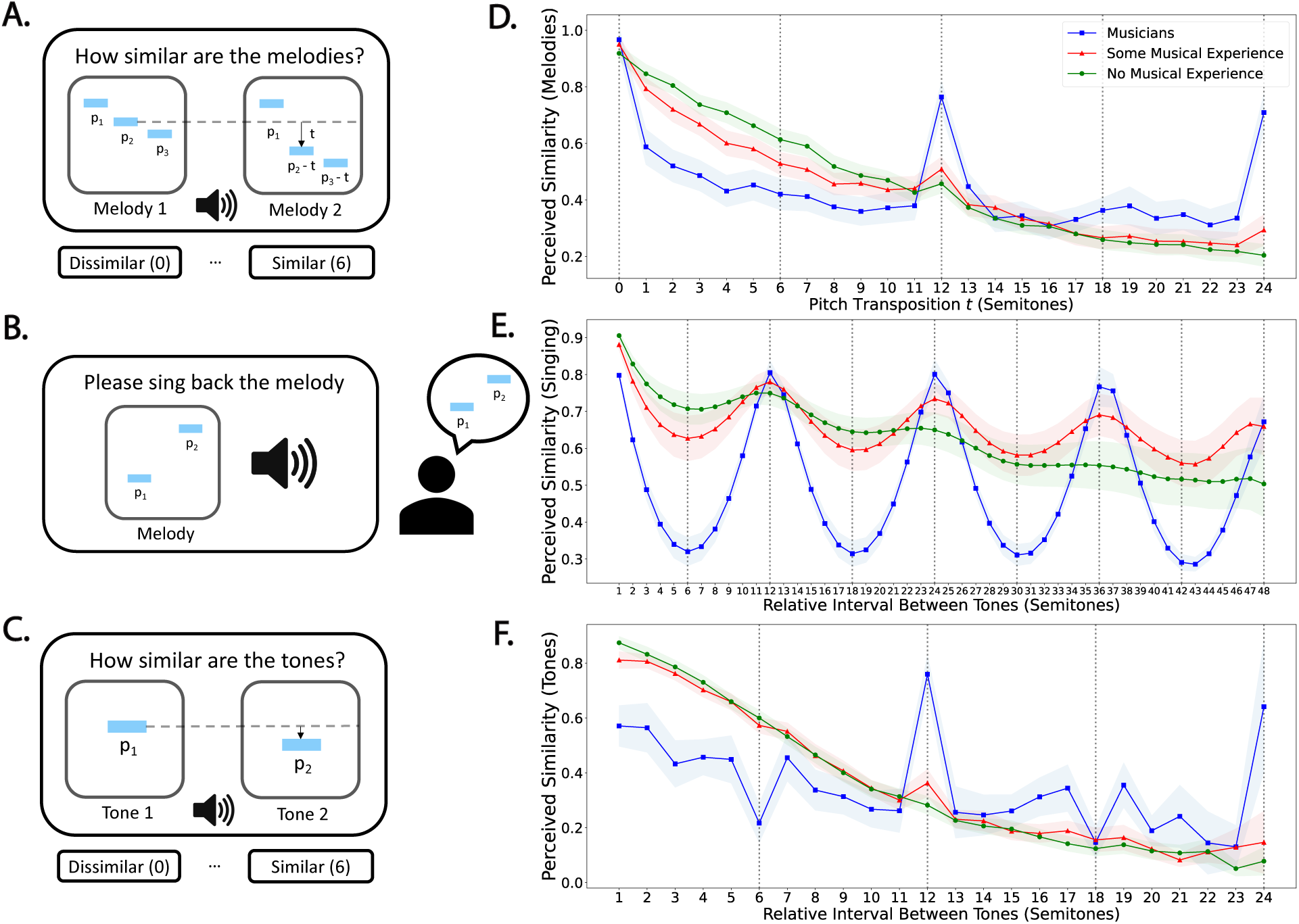
Probing pitch similarity across tasks and musical experience. Left panel: Schematics of the three paradigms **A.** similarity judgments over pairs of melodies that differ by a transposition, **B.** free imitation of two-note melodies through singing, and **C.** similarity judgments over pairs of isolated tones. Right panel (**D-F**): Corresponding normalized similarity profiles as a function of interval separations for the three behavioral paradigms and different participant groups (no musical experience, some musical experience, and musicians; see Methods). Shaded area, here and everywhere else, represents 95% confidence intervals bootstrapped over participants with 1,000 repetitions.

Crucially, we selected paradigms that allowed for a similarity-based analysis as with MDS and RSA (Shepard, 1980; Kriegeskorte et al., 2008). In these approaches, behavioral data such as direct similarity judgments, response distributions (e.g., confusion probabilities), or patterns of neural activation (Shepard, 1980; Kriegeskorte et al., 2008; Sucholutsky et al., 2023) are translated into similarity matrices (i.e., each matrix entry measures the similarity between a pair of stimuli) that can be used to study the global organization of a representation of interest. This is done by applying an embedding algorithm such as MDS (Shepard, 1980) which represents stimuli as points in a space and arranges them such that similar stimuli (based on the similarity matrix) are embedded close together, and dissimilar ones far apart. This results in a geometrical model of the psychological space without a priori assuming its structure. Accordingly, our experiments covered tones that spanned multiple octaves which allowed us to elicit similarity matrices across a broad range. Our goal here is to deploy the similarity-based approach to characterize the most suitable geometrical approximations of the structure of musical pitch and to test to what extent the helical model, or other models, provide a consistent and unified account across tasks and experience groups.

As for the specific paradigms, these were as follows. In the first paradigm, we wanted to probe pitch representations in the context of melody perception, which is one of the most prominent contexts that involve musical pitch perception across cultures (Savage et al., 2015; Mehr et al., 2019). To that end, we expanded the version of the Demany and Armand (1984) task used by Jacoby et al. (2019) whereby people listened to pairs of randomly generated three-note melodies and rated their similarity on a Likert scale (Figure 2A; see Methods). Crucially, we generated the second melody from the first by applying a fixed transposition *t* to the second and third tones across a wide two-octave range. Since these melodies were randomized across trials, we expected the average rating to be able to track the interplay between pitch height (how separated are the tones on a log-scale) and chroma (whether tone chroma is identical in both melodies, i.e., at octave transpositions).

In order to provide a complementary perspective on pitch representations, the second paradigm we considered was that of Jacoby et al. (2019) whereby people were asked to sing back two-note melodies that extended outside their singing range (Figure 2B; see Methods). This task provides an ecologically-valid form of musical engagement with pitch as singing is present in virtually every human culture (Savage et al., 2015; Mehr et al., 2019), and recent methodological advances enable it to be conducted online and at scale (Anglada-Tort et al., 2023). Moreover, singing also involves pitch production and as such complements the perceptual-evaluative task of melodic similarity presented above. It also allows one to derive an implicit measure of perceptual similarity based on behavior as participants are not asked to evaluate the similarity between any tone pairs. Participants heard two-note melodies, with the first tone sampled from a frequency range of 45.5 − 105.5 MIDI note corresponding to 113.2 − 3623.1 Hz, and were asked to reproduce them by singing (the participants’ singing range is approximately 80 − 1000 Hz). We specifically focused on how people approximated tones outside their singing range and constructed similarity scores between pitch values based on how similar their response distributions were (see Methods). This approach is consistent with prior multi-dimensional scaling analysis techniques whereby behavioral response distributions (e.g., confusion probabilities; Shepard, 1987) are used to construct similarity measures with the idea being that two stimuli are similar in so far as they produce the same behavioral response. This also enables us to analyze the production-based singing data using the same methodologies that we apply for the perceptual-evaluative similarity judgment paradigms.

For the final paradigm, we returned to the classic approach of Shepard (1980) and asked participants to directly rate the similarity between pairs of isolated tones with as little as possible additional context (Figure 2C; see Methods). We generated a high-powered similarity matrix by asking participants to rate pairs of harmonic complex tones taken from a two-octave range from C4 to C6.

As for participants, our goal was to cover a wide range of musical expertise levels. To do that, participants were recruited online from two pools (see Methods): Amazon Mechanical Turk (AMT) and an internal pool of professional musicians through the Max Planck Institute for Empirical Aesthetics (see Methods). Throughout, we divided the AMT participants into two categories based on their reported years of musical experience: those who reported zero years of musical experience (‘No Musical Experience’) and those who reported non-zero years of musical experience (‘Some Musical Experience’), in addition to the professional musicians category (‘Musicians’). All demographic information is provided in Supplementary Table A1.

Viewed together, our experimental design provides a holistic test for geometrical models of pitch across nine different combinations of expertise levels and tasks. We follow this analysis with a theoretical modeling section to further test the composite representation hypothesis, i.e., that the observed variation across conditions can be explained by the reweighting of a small set of perceptual factors. We next unpack the technical details of the studies in the Methods section, and then proceed to the Results followed by the modeling.

## Methods

### Software Implementation

The behavioral paradigms were designed and deployed using PsyNet^2^ (Harrison et al., 2020), a modern framework for complex experiment design which builds on the Dallinger^3^ platform for online experiment hosting and participant recruitment.

Participants interact with the experiment in the browser using a front-end interface which, in turn, communicates with a back-end Python cluster that organizes the experiment timeline. All experiment and analysis code are available in the OSF repository provided with this manuscript (see Transparency and Openness below).

### Stimuli

We defined absolute pitch using the Musical Instrument Digital Interface (MIDI) semitonal scale which is based on a logarithmic pitch representation. This was motivated by previous research which showed that participants across cultures use approximately logarithmic pitch representations (Jacoby et al., 2019). Specifically, the MIDI scale was defined as follows: *f* = 440 × 2^(^*^p−^*^69)^*^/^*^12^ where *f* is frequency in Hertz and *p* is the corresponding pitch value. This means that a Concert A (A4 or 440 Hz) corresponds to a value of 69 on this scale. All experiments involved additively synthesized harmonic complex tones 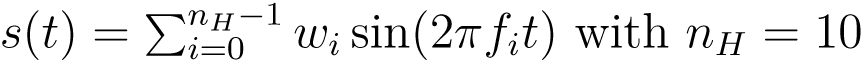 harmonics, where *f_i_* = *f*_0_ × (*i* + 1) for some fundamental frequency 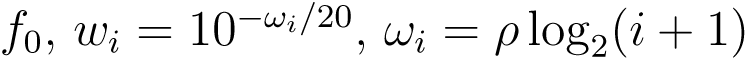 and a roll-off parameter, *ρ*, which parametrically represents the timbre of the sound by controlling the prominence of high harmonics. The tones were synthesized using Tone.js^4^, a Javascript library for audio synthesis in the browser. Each tone had a temporal envelope comprising of a linear attack portion, an exponential decay portion to some sustain amplitude, and a final exponential release portion. These were specified based on the constraints of each paradigm (see below).

### Similarity Paradigms

The similarity judgment paradigms used a roll-off parameter *ρ* of 3 dB/octave. In the similarity between tones paradigm, the envelope comprised a 200 ms linear attack segment reaching an amplitude of 1, followed by a short 100 ms exponential decay to an amplitude of 0.8, and finally an exponential release segment lasting for 1 second. The two tones within a given pair were separated by 300 ms of silence. In the case of melodies, the individual tones within melodies similarly had an envelope comprising of 200 ms linear attack followed by a short 100 ms decay to an amplitude of 0.8, and then an exponential release portion with the same decay rate as before. The release portion was played up to an inter-onset-interval of 650 ms in the first two tones (i.e. for 350 ms), and for 1 second in the last tone, followed by 200 ms of silence. This choice of parameters was intended to ensure that the presentation of melodies was brief and well-separated (so as not to interact with memory and to allow participants to experience a range of melodic pairs within a single experiment).

In the similarity judgment paradigm over melodies, random melodies were synthesized by first uniformly sampling a starting tone in the MIDI range 76-80, and then generating the second and third tones by subtracting a uniformly sampled integer interval in the range 5 − 7 and 9 − 11 semitones, respectively. This means that the resulting melodies could have a variety of melodic intervals per transposition so that pre-existing musical expectations would not prime the perception of the transposition interval.

Transpositions were then generated by subtracting a fixed integer interval in the range 0 − 24 semitones from the second and third tones. As for the similarity judgment paradigm over tones, tones covered the MIDI range 60 − 84 corresponding to the notes C4-C6.

### Singing Paradigm

In the singing task, the pitch range, timbre, and temporal envelope of the tones, can all influence the difficulty of singing back melodies. We thus adapted sound parameters that were extensively tested in a previous online singing study (Anglada-Tort et al., 2023) and used the following values: a roll-off parameter *ρ* of 14 dB/octave, a 10 ms attack, a 50 ms decay to an amplitude of 0.6, followed by a 550 ms release portion and 190 ms of silence (i.e., 800 ms inter-onset-interval between tones).

Two-note melodies were produced based on the design of Jacoby et al. (2019) (Experiment 4) by sampling the first tone from the range of 45.5 − 105.5, and then creating the second tone by adding one of the intervals 0, ±1, ±2, or ±3 semitones. To further assist singers in their responses we applied a method proposed in Jacoby et al. (2019), where each experiment consisted of a series of “blocks” presented in random order, each of which presented stimuli within a single frequency register (i.e., by sampling the first tone uniformly from the range [*p*_0_ − 6*, p*_0_ + 6] where *p_0_* = 51.5, 63.5, 75.5, 87.5, 99.5).

### Participants

Participants were recruited online from two participant pools: Amazon Mechanical Turk (AMT), and an internal pool of professional musicians through the Max Planck Institute for Empirical Aesthetics. To cover different experience levels, we divided the AMT participants into two subcategories based on whether they reported zero years of musical experience (‘No Musical Experience’) or some years of musical experience (‘Some Musical Experience’), in addition to the musicians (‘Musicians’). The division of the AMT population also aligns with a median-split based on reported years of musical experience. Participants reported years of musical experience by responding to the query: ‘Have you ever played a musical instrument? If yes, for how many years did you play?’. Demographic information and sample sizes for all participant groups are summarized in Supplementary Table A1. We confirmed in a post-hoc bootstrap analysis over participants that our sample sizes yielded reliable similarity data (average split-half data-data correlation *r_dd_* across conditions was.88; see Supplementary Table A3). Additional recruitment details are provided below (see also Appendix B for data-quality measures such as pre-screeners).

### AMT Participants

Participants on AMT were subject to the following recruitment criteria designed to ensure data quality: 1) participants must be 18 years of age or more, 2) they must reside in the United States, 3) they have a 99% approval rate or higher on prior AMT tasks, and 4) have successfully completed 5,000 tasks on AMT (a common threshold in large scale AMT studies; e.g., Hardy et al., 2023). These participants provided informed consent under a Princeton University Institutional Review Board (IRB) protocol (application 10859) in the similarity tasks and a Max Planck Ethics Council protocol (application 2021_42) in the singing task, and were paid a fair wage of 12 USD per hour.

### Musicians

The musician cohort was recruited through an internal online participant pool at the Max Planck Institute for Empirical Aesthetics (MPIEA). Participants were recruited to this pool by researchers and research assistants at MPIEA by sending emails to local and international music conservatories, as well as handing flyers at local music conservatories in Germany. The pool comprises the following countries: Germany, Spain, Italy, Malaysia, France, Hungary, United Kingdom, Switzerland, Australia, Peru, Indonesia, Argentina, Croatia, Hong Kong, Lebanon, Chile, and Serbia. In order to participate, musicians were required to be at least 18 years of age and to have at least 10 years of active musical training. Participants provided consent under a Max Planck Ethics Council protocol (application 2021_42) and were paid at a rate of 15 USD per hour. Participant took the experiment remotely, and the web-interface was identical to the one used by the AMT participants. We excluded three participants in melodic similarity, two in singing, and two in tone similarity who reported less than 10 years of musical experience.

We also note that the number of musicians somewhat varied between the similarity paradigms due to the nature of online recruitment whereby participants had the freedom to select which experiments they wanted to complete from a list of available studies.

### Procedure

#### Similarity Paradigms

In the similarity over melodies task, the experiment proceeded as follows: upon completing the consent form participants received the following instructions: “In this experiment we are studying how people perceive melodies. In each round you will be presented with two three-note melodies and your task will be to simply judge how similar they are. You will have seven response options, ranging from 0 (‘Completely Dissimilar’) to 6 (‘Completely Similar’). Choose the one you think is most appropriate. You will also have access to a replay button that will allow you to replay the sounds if needed. Note: no prior expertise is required to complete this task, just choose what you intuitively think is the right answer”. Participants then completed two practice trials and then proceeded to the main experiment. The procedure for the similarity over isolated tones task was identical, except that we replaced “melodies” in the instructions with “sounds”. Participants completed up to 80 trials.

#### Singing Paradigm

After providing informed consent and completing the pre-screening trials (see Pre-screening in Appendix B) where they were also familiarized with the singing task, participants received the following instructions: ‘In this final part of the experiment, we will ask you to do exactly the same as in the previous part. 1. You will hear a 2 note melody. 2. You will have to sing back the 2 notes. That means, in total you will sing 2 notes. The difference is that this time there will be a total of 38 trials. For all notes, use the syllable TA to sing the note and leave a silent gap between notes. Note: some of the tones you’ll hear will be high, and well above your singing range. You should, however, sing them within your comfortable singing range.’ The latter note was intended to ensure that participants were not surprised by the high tones (consistent with Jacoby et al. (2019); see Session Structure in that paper), as well as to ensure that they imitated the melodies within their natural singing range. Participants then completed the trials which corresponded to the randomized trial blocks (see Stimuli) in addition to three repeat trials (i.e. repeating three random trials from the original 35). The trials were presented in one continuous stream without any additional indications. In a given trial, participants received the following instructions: “This melody has 2 notes. Sing back the melody. Use the syllable TA”. The recording interface comprised of a progress bar (see Supplementary Figure A9) indicating the three recording stages. The initial portion, indicated in orange, signaled the listening portion (“Listen to the melody…”). Then, a second portion, indicated in red, signaled that the recording was in progress (“Recording…SING THE MELODY!”). This portion lasted for the duration of the original melody plus one second. Finally, a third portion, highlighted in green, indicated that recording was complete (“Done!”), and lasted for 500 ms. The next trial then started automatically, and so on.

## Data Analysis

### Constructing Similarity Matrices

To generate an aggregate similarity matrix from the direct similarity judgment paradigms, we applied the following procedures. In the case of similarity over isolated tone pairs, we simply computed the average Likert score per pair of items and divided by a constant of 6 so that similarity scores would be normalized between 0 and 1 (rather than 0 and 6). As for similarity over melodies, since the data is one-dimensional in that case (a pitch transposition *t* between random melodies) we constructed a two-dimensional matrix *a_ij_* from the one-dimensional data *s_t_* using the formula *a_ij_* = *s_|pi−pj |_* where *p_i_* and *p_j_* are pitch values of interest. We again divided by a factor of 6 so that similarity scores would be normalized between 0 and 1. As for the singing data, the procedure was slightly more complicated and is described in Appendix B (see Singing Response Distributions). This procedure was inspired by representational similarity analysis (RSA; Kriegeskorte et al., 2008) where the similarity of stimuli is defined by the similarity of their corresponding behavioral or neural responses. We confirmed in a simulation analysis that this approach is flexible enough to capture different behavioral patterns and structures (see Appendix B and Supplementary Figure A5). Throughout, we complement our approach with additional analyses of the raw data (e.g., marginals and response distributions; see below). To produce smoothed raw matrices, we averaged the similarity matrices over their diagonal values, i.e.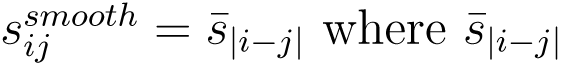 is the average over the |*i* − *j*|-th off-diagonal of *s_ij_* (0 being the main diagonal).

### Multi-dimensional Scaling

We constructed MDS embeddings using the MDS method of scikit-learn. This was done in two steps, first we applied a metric MDS to find an initial embedding which was then fed as an initialization into a non-metric MDS. We used three components, a maximum iteration value of 10,000, and a tolerance parameter of 1e-100. We chose three dimensional embeddings based on a stress curve analysis (Supplementary Figure A2) and selected the smallest number of dimensions for which all stress values across all conditions dropped below 0.15 (MDS fit is deemed poor above this threshold; Kruskal, 1964). This ensured that all structures are represented adequately within the same unified presentation (since lower dimensional structures can always be represented in higher dimensions but not vice versa). Finally, we note that all MDS solutions presented in this work are also available as 3D interactive HTML visualizations available through the OSF repository (see Transparency and Openness).

### Transparency and Openness

Data and code used in this work can be accessed via the link: https://osf.io/nx586/?view_only=a0f74c9269a4448f821aadccf76ba175 (this link will be replaced with a permanent link upon publication). This study was not preregistered. We report how we determined our sample size, all data exclusions, all manipulations, and all measures in the study.

## Results

### Empirical Similarity Profiles

Starting from the first paradigm, recall that participants rated the similarity between pairs of randomly generated three-note melodies where the second melody was produced from the first by applying a fixed transposition *t* to the second and third tones across a wide two-octave range. Figure 2D shows substantial group differences in the average profile as a function of the transposition interval (see Participants and Table A1 for all demographic and experience information). Participants with no musical experience (*N* = 102; “No Musical Experience” in Figure 2), exhibited a predominantly linear profile (linear model explains 93.8% of the variance of the average profile, CI: [91.8, 95.8]) with a small but significant bump at the first octave (12 semitones, CI of the mean rating difference between 12 and the average rating of 11 and 13 semitones is [0.030, 0.084] which does not include zero), and musicians (*N* = 41) exhibiting a highly non-linear profile (linear model explains only 17.4% of the variance, CI: [7.98, 26.9]) that is more flattened with significant spikes at octave transpositions (12 and 24 semitones; CIs for mean rating difference for 12 and the average of 11 and 13, and 24 and 23 semitones were [0.293, 0.406] and [0.307, 0.440], respectively) suggesting octave equivalence and chroma sensitivity.

Participants with some musical experience (*N* = 92; “Some Musical Experience”), on the other hand, exhibited a profile that is somewhere between those of the other two groups. Quantitatively, a linear model explained 86.0% of the variance of the average profile (CI: [80.7, 91.3]) and the CI of the mean difference in rating between 12 and the average of 11 and 13 semitones, and between 24 and 23 semitones were [0.061, 0.131] and [0.016, 0.088], respectively.

Moving to the second paradigm, here we analyzed how participants imitated two-note melodies that are outside their singing range by evaluating their response distributions (see Methods). We estimated similarity scores between different target pitch values by computing the Jensen-Shannon distance (JSD) between their response distributions (i.e., the distributions of pitch values produced when participants attempted to imitate them). This was done by first applying a two-dimensional kernel density estimation to target-response pitch pairs and then applying JSD between slices pertaining to pitch values of interest (see Methods; raw target-response distributions are provided in Supplementary Figure A1). Figure 2E shows the resulting average similarity profiles.

Similar to the first paradigm, different experience groups exhibited qualitatively different profiles, with participants with no reported musical experience (*N* = 50; see Table A1 for full demographics) showing a linear similarity trend with weak residual periodicity at the octave (linear model explains 89.0% of the variance with CI [78.6, 99.4], and a sinusoidal model with 12-semitone periodicity explains 7.4% of the variance with CI [0.6, 14.3]) and musicians (*N* = 41) exhibiting a highly periodic pattern with strong peaks at integer multiples of the octave (linear model explains 1.2% of the variance with CI [0, 3.3] and a sinusoidal model with 12-semitone periodicity explains 92.7% of the variance with CI [89.8, 95.6]). Participants with some musical experience (*N* = 49) yielded a pattern that is in between the two other profiles (linear model explains 28.5% of the variance with CI [0.2, 56.8] and a sinusoidal model with 12-semitone periodicity explains 61.0% of the variance with CI [32.8, 89.3]).

Finally, we turn to the third paradigm which involved participants directly rating the similarity between tone pairs that varied in the range C4 - C6. Figure 2F shows the average similarity rating as a function of the interval between the two tones. In this case, participants with zero reported years of musical experience (*N* = 94; additional details in Table A1) exhibited a nearly perfectly flat profile (linear model explains 90.8% of the variance with CI [89.1, 92.6]) with no significant peak at the octave: CI of the mean rating difference between 12 and the average rating of 11 and 13 semitones is [−0.02, 0.042]), and musicians (*N* = 30) on the other hand exhibiting a highly non-linear pattern (linear model explains 18.2% of the variance with CI [1.4, 35.0]) with spikes at the octaves (12 and 24 semitones; CIs of the mean difference in rating between 12 and average of 11 and 13 semitones, and between 24 and 23 semitones were [0.421, 0.580] and [0.240, 0.780], respectively) and sharp dips at the tritones (6 and 18 semitones; CIs of the mean difference in rating between 6 and average of 5 and 7 semitones, and between 18 and average of 17 and 19 semitones were [−0.315, −0.155] and [−0.276, −0.130], respectively) indicating strong octave equivalence but also strong tritone aversion and enhanced similarity at perfect intervals around it (i.e., 5, 7, 17 and 19 semitones). Participants with some musical experience (*N* = 92) capture the onset of the transition between the prior two groups (linear model explains 89.5% of the variance with CI [85.6, 93.4]) and a small but significant peak at the first octave (CI of mean difference between 12 and the average of 11 and 13 semitones was [0.047, 0.148]) and an onset of a dip at the first tritone (CI of mean difference between 6 and the average of 5 and 7 semitones was [−0.058, −0.006]), but not around 18 semitones (CI of mean difference between 18 and the average of 17 and 19 semitones was [−0.063, 0.021]).

### Derived Geometrical Models

The observed similarity marginals suggest strong task and experience dependencies.

To see what geometrical models could underlie those marginals and whether they are consistent with a unified pitch helix, we subjected them to three-dimensional MDS analysis (we selected three dimensions based on a stress curve analysis; see Supplementary Figure A2). This corresponds to averaging the raw similarity matrices over their diagonals (i.e., over equal semitonal separations; see Supplementary Figures A3-A4) and applying MDS (see Methods). The resulting MDS solutions, shown in Figure 3, exhibit a rich array of geometries, with the traditional helical representation appearing in only three out of the nine conditions considered (Figure 3D-F) and pertaining to the singing task (we confirmed in a simulation analysis that this is not an artifact of the singing paradigm and that it is perfectly capable of capturing other structures, see Supplementary Figure A5). In addition, MDS solutions for the raw unprocessed similarity matrices exhibit similar qualitative differences, though they are naturally noisier (Supplementary Figure A6). As with Figure 2, we see that with increased musical experience the derived representations become increasingly non-linear. In the melodic similarity case, participants with no musical experience yield a largely linear solution (Figure 3A), whereas musicians yield a solution characterized by octave-equivalent tone units (Figure 3C). Participants with some musical experience yield an intermediate non-linear solution (Figure 3B). As for the singing paradigm (Figure 3D-F), while all groups yield helical solutions, those solutions get increasingly tighter with experience, possibly reflecting an increasingly diminished pitch height dimension. Finally, for the isolated tone similarity paradigm, we also observe a structural transition as a function of experience whereby the linear approximation observed in the lower experience groups (Figure 3G-H) factorizes into a double helix in musicians (Figure 3I) to account for tritone aversion and heightened similarity at perfect intervals.

**Figure 3.**
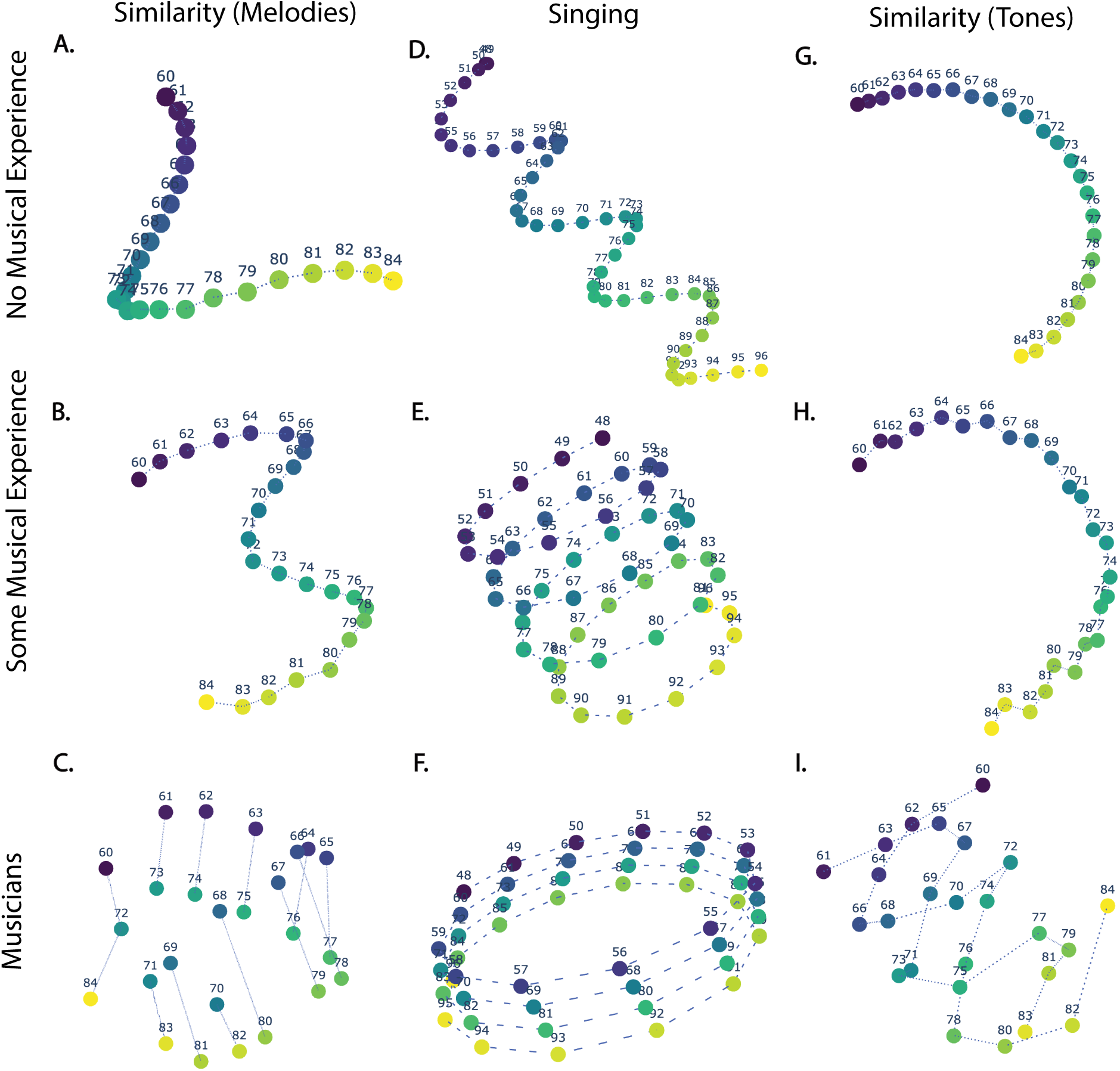
Multidimensional scaling solutions in three dimensions for the various similarity data shown in Figure 2. Left to right: similarity judgments over melodies that differ by a transposition (**A-C**), free imitation of two-note melodies via singing (**D-F**), and similarity judgments over pairs of isolated tones (**G-I**). Numbers and colors indicate pitch values in MIDI scale. Interactive 3D versions are available in the OSF repository for all MDS solutions shown in the present work (see Transparency and Openness).

Curiously, this recovers the ‘double-helix wound around a line’ structure hypothesized by Shepard (1982a, 1982b) as a candidate approximation for the structure of musical pitch (Supplementary Figure A12B). It differs, however, from the four dimensional double helix solution derived by using the probe tone data of Krumhansl and Shepard (1979) which is wound around a torus (the direct sum of the chroma circle and the circle of fifths; Figures 3 and 5 in Shepard, 1982a; see also Supplementary Figure A12C). Viewed together, these results provide clear evidence that a simple helical model, or in fact any fixed geometrical model, is inadequate for capturing musical pitch perception across tasks and experience.

### Computational Modeling

The results so far suggest that different task and musical experience conditions load differently on pitch-related psychological attributes. To test whether the observed variations in similarity patterns can be captured by a configurable basis of perceptual factors, we used a computational modeling approach inspired by an analysis initially proposed by Shepard (1982b) and adapted here to capture the different possible sources of variance observed in our data. Let *s_ij_* denote a similarity matrix that encodes the perceived similarity between stimuli *p_i_* and *p_j_* (without loss of generality we assume that 0 ≤ *s_ij_*≤ 1), and let Δ*_ij_* = 1 − *s_ij_* denote its corresponding dissimilarity matrix. Our goal is to find a decomposition of Δ*_ij_* in terms of a parametric basis of dissimilarity measures or “components” *d^(c)^_ij_*, each corresponding to a different perceptual factor (Figure 4; see Modeling Methods below). Mathematically, we seek a solution of the form

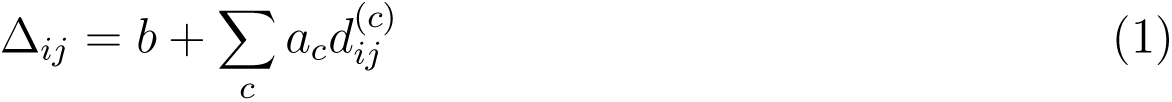

**Figure 4.**
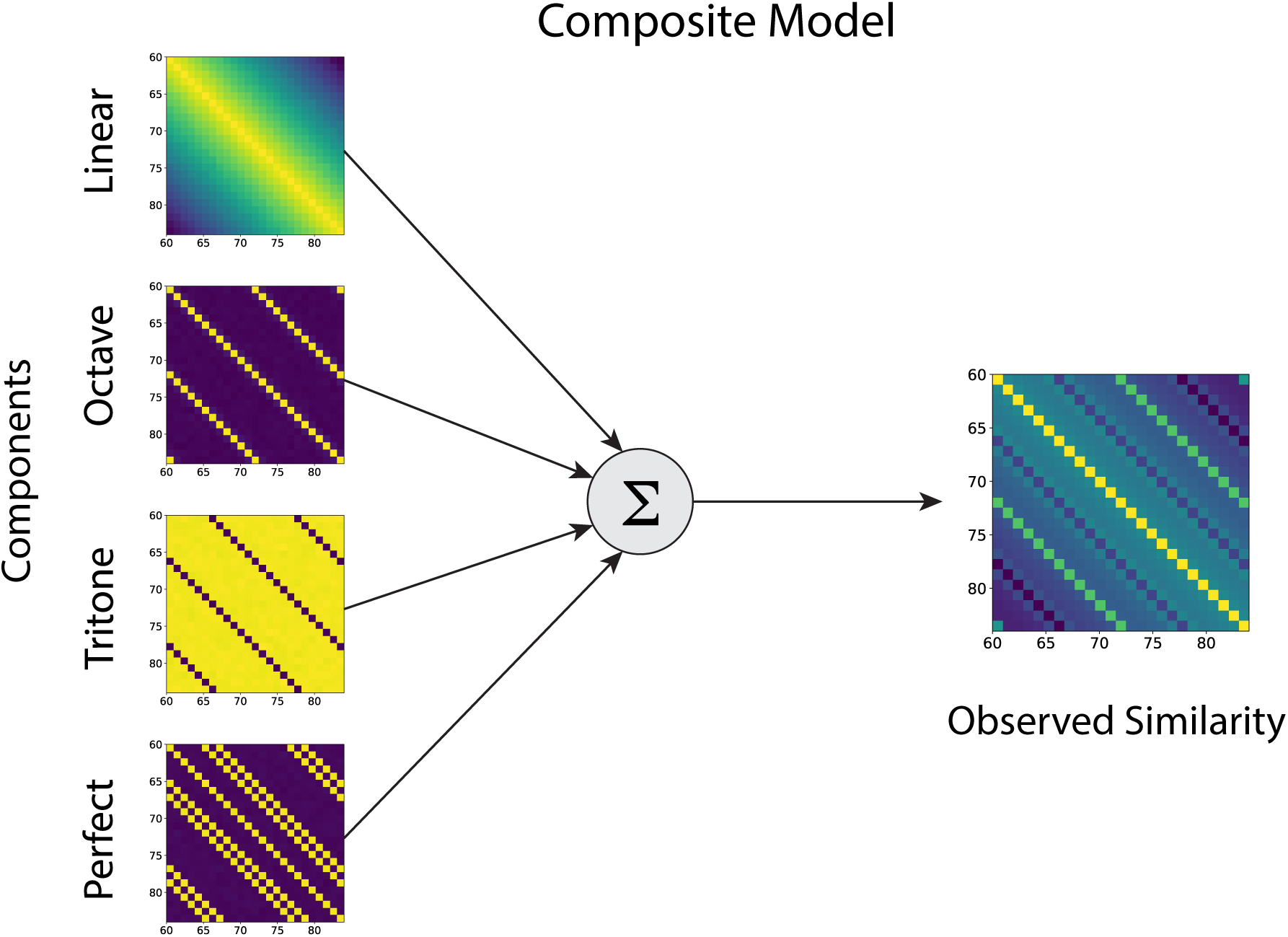
Composite model schematic. Distinct perceptual components (log-linear height, octave equivalence, tritone aversion, perfect interval equivalence) are combined to produce an observed similarity matrix.

**Figure 5.**
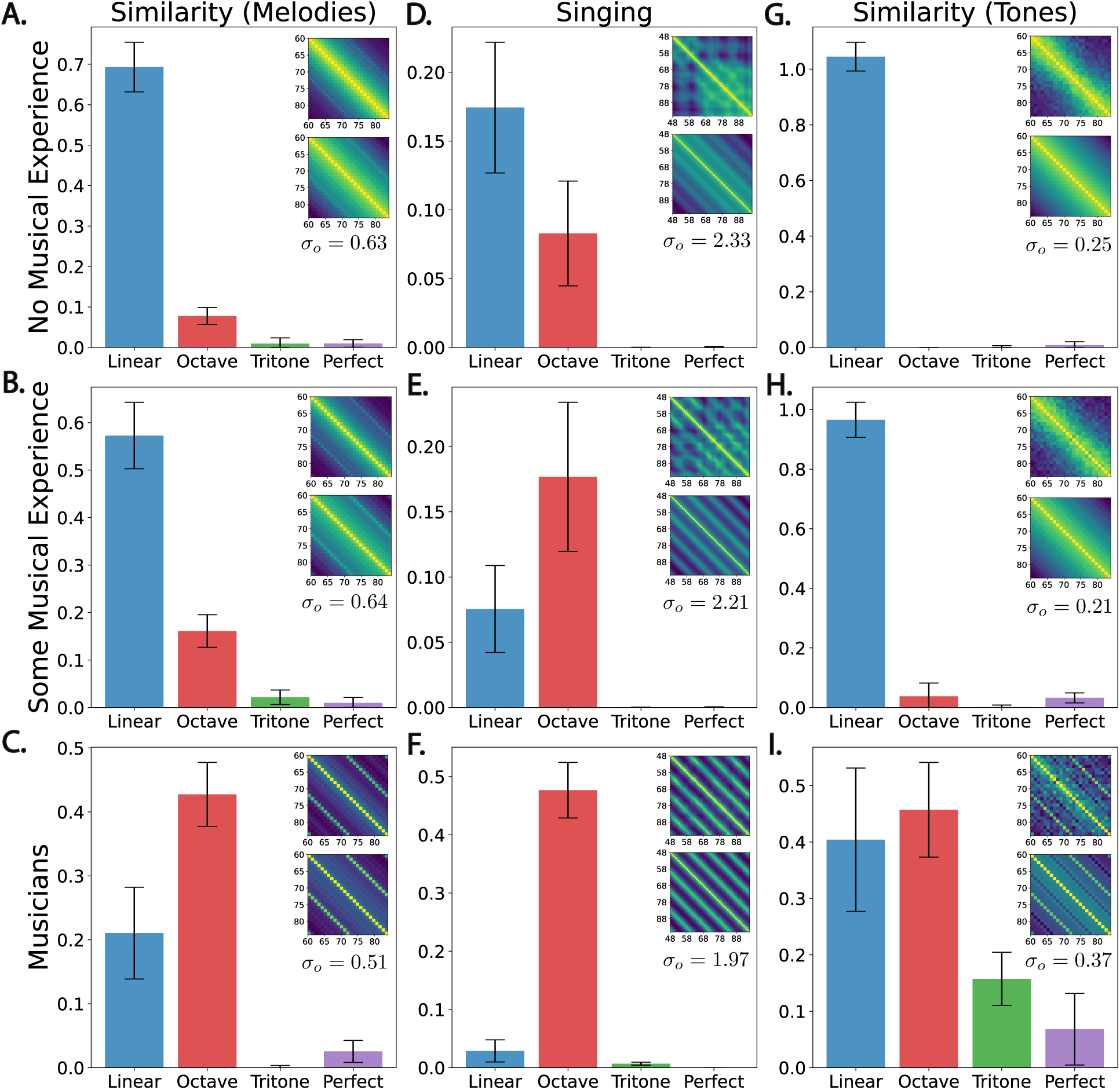
Perceptual component analysis of similarity. Fitted component coefficients to data from different musical experience groups and their associated raw (top) and model (bottom) similarity matrices shown as insets (enlarged insets are shown in Supplementary Figures A3 and A7; numerical values are provided in Supplementary Table A2). Error bars indicate 95% confidence intervals bootstrapped over participants with 1,000 repetitions.

where *b, a_c_* ≥ 0 are non-negative coefficients. Based on the marginal analysis of Figure 2, we selected four components: a linear pitch height scale *d_ij_^l^*, and three additional components that represent the identification of particular intervals, namely, a component d_ij_*^o^* that identifies octaves, a component *d_ij_^p^* that identifies perfect intervals (fourth and fifth), and a component *d_ij_^t^* that captures aversion to tritones. The octave component had an additional width hyperparameter *σ_o_* that interpolated between sharp octave recognition and smoother chroma matching (see Modeling Methods for explicit formulae).

## Modeling Methods

### Perceptual Components

The explicit formulae for the different components were: a (log) linear pitch scale *d_ij_*^(^*^l^*^)^ ∝ |*p_i_* − *p_j_*|, two positive interval recognition components *d^(c)^_ij_* = 1 − exp (−(*χ*(*p_i_, p_j_*) − *c*) */*2*σ^2^_c_*) where *χ*(*p_i_, p_j_*) = min{*ψ_ij_,* 6 − *ψ_ij_*} and *ψ_ij_* = |*p_i_* − *p_j_*| mod 12 to capture chroma distance (i.e., intervallic distance irrespective of register) and *c* = 0, 5 for octave (*d*^(0)^*_ij_*) fifth; *d^(p)^_ij_*), respectively, and one negative recognition component *d_ij_*^(^*^c^*^)^ = exp (−(*χ*(*p_i_, p_j_*) − *c*)^2^*/*2*σ_c_*^2^) with *c* = 6 to capture tritone aversion (*d_ij_*^(^*^t^*^)^). We fixed *σ_c_* = 0.25 semitones for the tritone and perfect intervals as a narrow interval recognition threshold and optimized the octave width *σ_o_* as a hyperparameter since different values of *σ_o_* efficiently interpolate between sharp octave recognition and smooth distance on the chroma circle (since *c* = 0). As a sanity check for the chosen width parameter values, we refitted the model in the musician tone similarity condition (Figure 5F, the only condition in which the tritone and perfect intervals had substantial contributions) but this time allowing all width parameters to vary. We found that the optimal width parameters did not differ much from our chosen values (*σ_o_* = 0.39, *σ_t_* = 0.17 and *σ_p_* = 0.57). In addition, to ensure that the coefficients are properly normalized (i.e., each component varied between 0 to 1), we rescaled the linear component such that its largest separation on a given range of interest equals to one (e.g. if pitch differences varied between 0 and 24 semitones, we defined *d_ij_*^(^*^l^*^)^ = |*p_i_* − *p_j_*|*/*24).

### Model Fitting and Evaluation

We fitted components to the data using the LinearRegression method of scikit-learn. This was done by first flattening the upper triangular part of each component’s distance matrix *^d^ij*^(^*^c^*^)^ into a feature vector *vi*^(^*^c^*^)^ and then fitting a linear regression of the form 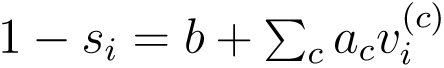 where *b, a_c_* ≥ 0 are non-negative coefficients and *s_i_* is the flattened upper-triangular matrix of the behavioral similarity data.

We repeated this fitting process 1,000 times by bootstrapping over participants in a split-half fashion (i.e., generating random half-splits of the data based on participants and then fitting the model to one half and testing on the other). The optimal octave width hyperparameter *σ_o_*was fine-tuned using the blackbox optimizer scipy.optimize.minimize over the bootstrapped linear regression procedure described above. To quantify model performance, we computed three Pearson correlation coefficients for each of the 1,000 split-halves as follows: *r_dd_* the correlation between the corresponding human similarity matrices of each split, *r_dm_* the correlation between the model fit on one half with the human similarity of the other (there are two ways to compute this so we took the mean), and *r_mm_* the correlation between the fitted models on each half. We then used these values to compute the corrected correlation for attenuation 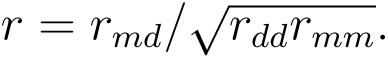

## Modeling Results

Despite the economy of the component model, it provided excellent fit to the observed data with mean Pearson correlation of *r* =.94 (*r* values ranged between.80 −.99 depending on the experiment and were corrected for attenuation; see Modeling Methods for details regarding model evaluation; full list of correlations and other evaluation metrics is provided in Supplementary Table A3).

The resulting component weights for each of the paradigms are summarized in Figure 5 (the numerical values for all parameters can be found in Supplementary Table A2; enlarged insets are provided in Supplementary Figures A3 and A7). We see that these weights vary widely depending on experience and task. Starting from the melodic similarity paradigm (Figure 5A-C) we observe that most of the variability is driven by the linear and octave components for all participant groups (Musicians: *r* =.98, 95% CI: [.98,.98]; Some Musical Experience: *r* =.98, 95% CI: [.97,.98]; No Experience: *r* =.98, 95% CI: [.98,.98]), however the octave component was significantly more dominant for musicians (Figure 5C; octave coefficient CI [0.38, 0.48]) compared with the lower experience groups (Figure 5A-B; octave coefficient CI [0.13, 0.19], and [0.06, 0.10] for participants with some musical experience, and no musical experience, respectively). Moreover, the relatively diminished linear component in musicians (Figure 5C; compare linear coefficient CI [0.14, 0.28] vs. octave coefficient CI [0.38, 0.48]) as well as the narrow octave width (*σ_o_* = 0.51) are suggestive of sharp octave recognition, whereby participants recognize the interval of an octave specifically as more similar without this enhanced similarity influencing other neighboring intervals (consistent with the flattened profile in Figure 2D).

Turning next to the singing paradigm (Figure 5D-F), we see an analogous pattern where the data are largely explained by the linear and octave components (Musicians: *r* =.98, 95% CI: [.98,.99]; Some Musical Experience: *r* =.84, 95% CI: [.80,.88]; No Musical Experience: *r* =.80, 95% CI: [.77,.83]). In particular, we see that the linear component in musicians is nearly absent relative to that of the octave (compare linear component CI [0.01, 0.05] vs. octave component CI [0.43, 0.53]) which is suggestive of chroma matching. Unlike the previous paradigm, however, here we see that the optimal octave component width is much larger (*σ_o_* = 1.97). Possible reasons for this result could be a perceptually broad representation (as a result of the higher task demands), and production noise. Interestingly, the reversal in the dominance of the linear and octave components in musicians relative to the participant group with no reported musical experience (Figure 5D) seems to happen also for the intermediate group with some musical experience (Figure 5E; linear component CI [0.05, 0.11] vs. octave coefficient CI [0.12, 0.24]) which is not the case for the melodic similarity paradigm (Figure 5B; linear component CI [0.50, 0.64] vs. octave component CI [0.13, 0.19]), suggesting that for non-musicians pitch chroma is more salient in the singing task than in the melodic similarity task (even though the singing task is perfectly capable of detecting a linear representation; see simulation results in Supplementary Figure A5).

Finally, for the similarity over isolated tones paradigm (Figure 5G-I) we see a distinct pattern in which the lower experience groups (Figure 5G-H; No Musical Experience: *r* =.96, 95% CI: [.96,.96]; Some Musical Experience: *r* =.97, 95% CI: [.96,.97]) load almost exclusively on the linear component, while musicians (*r* =.93, 95% CI: [.89,.97]), on the other hand, exhibit a complex pattern in which all components contribute, in particular, tritone aversion (coefficient CI: [0.11, 0.21]) and enhanced similarity at perfect intervals (coefficient CI: [0.01, 0.13]), in addition to relatively balanced linear and octave components (linear coefficient CI: [0.27, 0.53], octave coefficient CI: [0.38, 0.54]).

## Discussion

Our results provide strong evidence that the psychological representation of musical pitch cannot be adequately captured by a simple helical model as originally proposed by Shepard and commonly depicted in textbooks. On the one hand, this model exaggerates the effect of octave equivalence in non-musicians, and on the other, it misses the implications of alternative sources of variance in the judgments of musicians, namely, those pertaining to tritone aversion and enhanced similarity at perfect intervals. Moreover, our results show that while more complex geometrical models such as Shepard’s double helix may be useful for capturing musicians’ judgments in some tasks (similarity over tones), this may not be true for others (similarity over melodies and singing). Instead, our work highlights that the most suitable geometrical approximation is task dependent, reflecting a representation that is shaped by different psychological factors with variable degrees of salience which we model. Surprisingly, we also found that octave equivalence was most salient across experience groups in the singing task, despite being the most demanding and prone to production noise.

### Musical Pitch Perception is Composite and Task Dependent

Across all experiments, we found that four components explained most of the meaningful variance in pitch similarity (an average raw *R*^2^ of.79 across datasets, and an average corrected Pearson correlation of *r* =.94). However, their relative importance varied greatly between populations and tasks. Finding population and task dependency in pitch perception supports the idea that pitch is processed in higher-order brain areas (Nelken, 2011; Schnupp et al., 2011).

We found that a simple linear component dominated the responses of non-musicians, and also contributed to the responses of musicians (Figure 5). This is in line with the widely adopted definition of pitch by the American National Standards Institute (ANSI, 1960; “pitch is the auditory attribute of sound that allows sounds to be ordered on a scale from low to high”) which emphasizes the linear nature of pitch. This is also consistent with Jacoby et al. (2019) who showed evidence that log-linear pitch perception is present in participants across cultures.

The other leading component we found was octave equivalence. Our findings support the idea that octave equivalence varies across participants and that the extent to which it is manifested is shaped by its salience in the task, supporting Jacoby et al. (2019), Pressnitzer and Demany (2019), and Regev et al. (2019). More specifically, we show quantitatively how the strength of octave equivalence varies with musical experience within the same task (Figure 5). Moreover, we found that the strength of the octave component relative to the linear component was particularly salient in the singing task. This is surprising as one would expect the singing task to yield noisy response patterns being a production-based task and more challenging. While we did find that participant responses were noisy (Supplementary Figure A1), they nevertheless showed the most clear octave equivalence compared with other tasks, even though the singing task is capable of capturing a linear representation (Supplementary Figure A5A-B). This underscores the importance of naturalistic paradigms when evaluating pitch perception.

Beyond these two components our results showed that musicians in the similarity judgment task over tones also relied on two other components, specifically “aversion” to tritones and enhanced similarity at perfect intervals. What can explain the behavioral differences in musicians’ data across tasks? One idea is that musicians can identify musical intervals based on their ear training, and then report what they think they are expected to report based on their explicit knowledge of Western music theory (Aldwell et al., 2018), or reflecting their internalized known distribution of melodic intervals in musical corpora (Vos & Troost, 1989; Zivic et al., 2013; Anglada-Tort et al., 2023). However, the fact that we observed task dependencies for all three experience groups, including groups with zero years of musical experience, suggests that the results may not be fully attributed to participants “rationalizing” about their expected judgments. This is also consistent with the fact that in the singing task the derived pairwise similarity measure was implicit since participants did not directly provide a similarity judgment. Another possibility is that other aspects of music perception may influence musicians’ responses. For example, the phenomenon of melodic consonance, or the perceived pleasantness of tone sequences.

Recent research suggests that Western participants (Anglada-Tort et al., 2023) exhibit a hierarchy of preferences when evaluating the pleasantness of two-tone melodies, with tones separated by an octave and perfect intervals being particularly pleasant, and those separated by a tritone being particularly unpleasant. This pattern also overlaps to a certain extent with that of harmonic consonance (i.e., the pleasantness of simultaneous tones; Trainor and Heinmiller, 1998; McDermott et al., 2010; Harrison and Pearce, 2020; Lahdelma and Eerola, 2020; Marjieh, Harrison, et al., 2024).

Overall, our results align with findings from previous smaller-scale, indirect studies and consolidate them within a single, unified large-scale study that explicitly addresses the question of integrated geometrical models of musical pitch. Furthermore, our paper explicitly introduces the dimensions of task and expertise dependency (which are often studied separately) by evaluating multiple tasks in parallel across multiple levels of expertise, and verifying their joint influence on pitch perception.

## Implications

One possible implication of our findings is that distinct elements such as octave equivalence and logarithmic frequency scaling are perceived and processed separately by the brain. When individuals make judgments—such as assessing the similarity between two tones—they weigh these different features in a task-specific manner. When performing another task, such as singing to reproduce a tone, they may adjust their weighting of available cues or features differently. Previous research has shown that local pitch perception is task-dependent, and has leveraged this dependency to uncover distinct mechanisms for processing sounds into pitch. Indeed, by analogy to those psychoacoustic studies (Demany & Ramos, 2005; McPherson & McDermott, 2018), one may expect that the relative salience of different musical dimensions underlying the global organization of pitch percepts would be heavily modulated in different tasks, which is what we indeed find.

Our results carry implications for studying the neural correlates of pitch. Instead of a single representation in localized areas, we now expect pitch cues to be processed separately in lower and mid-level regions (Schulze et al., 2002; Bendor & Wang, 2005; Schnupp et al., 2011; Norman-Haignere et al., 2013; Feng & Wang, 2017), with integration only occurring at later stages of auditory processing (Nelken, 2011), akin to phoneme processing in speech (Mesgarani et al., 2014). Furthermore, our findings underscore the role of statistical learning (Loui et al., 2010; Loui, 2012; Pearce, 2018), as the salience of these components strongly depends on musical expertise (Schön & François, 2011; Jacoby & Ahissar, 2013). Our results also highlight the capacity of brain plasticity to strongly modulate auditory perception, consistent with a large body of work that shows differences between musicians and non-musicians in auditory perception and production, as well as their neural correlates (Shepard, 1982a; Koelsch et al., 1999; Gaser & Schlaug, 2003; Zatorre et al., 2007; Dellacherie et al., 2011; François et al., 2013; Jacoby & Ahissar, 2013; Merzenich et al., 2013).

Beyond neuroscience, our results are valuable for computational models of pitch perception. Predictive models of human behavior should consider both task dependency and expertise, as these are crucial for accurately capturing the nuances of pitch perception as demonstrated in the present work. Computational models that rely on mechanisms that cannot capture task and expertise dependencies should therefore be revised or excluded.

Likewise, future pedagogical resources (e.g., textbooks) depicting pitch representations should acknowledge that the pitch helix is only one instance of a broader class of geometrical approximations of pitch representations, some of which may be more suitable depending on the task and expertise considered.

For cognitive science, our findings indicate that pitch representations are composite rather than integrated, arising from the task-dependent influence of multiple, though limited in number, cognitive factors. This insight has broader implications, particularly when investigating perceptual representations in other modalities which are often presumed to be integrated. Moreover, recent research comparing human and machine learning representations commonly assumes integrated human representations to achieve alignment (e.g., Sucholutsky et al., 2023). Our results highlight the necessity of further considering composite, rather than purely integrated, representations when addressing human-machine alignment across a generalizable set of tasks.

### Limitations and Future Work

We end by discussing limitations which point to important directions for future research. First, our participant cohort is largely Western, particularly from the United States and Germany. This limits the generalizability of the present findings as elements of pitch perception such as octave equivalence and melodic preferences vary cross-culturally (Jacoby et al., 2019). A natural follow-up, therefore, could look at the way representations vary across cultures by applying the same methods deployed in this work which by design are suitable for cross-cultural research (similar to McDermott et al., 2016; Lahdelma and Eerola, 2020; McPherson et al., 2020; Jakubowski et al., 2022; Jacoby et al., 2024; Ozaki et al., 2024).

Second, in the present work we exclusively focused on population-level analysis of representations, however, since musical experience is very subjective one might expect to see individual differences (McDermott et al., 2010) and use that to inform the relation between components beyond group differences. Since we sought to derive dense similarity matrices that spanned multiple octave ranges (which necessitate hundreds of judgments to cover once; e.g. covering all pairs in the octave range C4-C6 requires at least 25 × 24*/*2 = 300 without repetition) while ensuring that the online studies are not too long to maintain high data quality, we were unable to collect enough data per participant to enable individual-level perceptual component analysis. Multi-session in-lab studies using the same paradigms of the present work may be a suitable follow up. This is particularly important, since splitting the data by musical expertise does show significant differences even within the same paradigm, this indicates that individual variation is likely to play a significant role in the structure of pitch representations (Figures 2, 3, and 5).

Third, previous work suggests that tonality may also be a shaping force in pitch perception (Krumhansl, 1979; Krumhansl & Shepard, 1979; Fogel et al., 2015). While our method tried to minimize carry-on effects of tonality from previous trials (by shifting tones randomly from trial to trial), it is possible that some sense of tonality can be induced even within a single trial, at least when it comes to expert musicians (Figure 5I).

Finally, there are other established paradigms for studying pitch representations that exist in the literature which we did not consider (e.g., Hoeschele et al., 2012) as they were not compatible with a similarity analysis. Future work could explore alternative analytical approaches to examine these cases and assess if similar patterns emerge. We intend to pursue these directions in upcoming research.

To conclude, while pitch perception provides the fundamental backbone underlying both speech and music perception, it is neither simple nor static. Instead of a unified helical representation, our results reveal a highly complex and dynamical phenomenon that is likely to depend on multiple independent factors. More broadly, our work showcases how combining large-scale behavioral studies with diverse behavioral paradigms and computational modeling can provide new answers to fundamental questions in auditory research. We believe that scaling up psychological research in tandem with concurrent computational approaches will continue to provide exciting new hypotheses concerning the nature of human cognition.

## Author Contributions

All authors contributed to the design of the experiments.

Author RM implemented the experiments, analyzed the results, and drafted the initial manuscript. All authors collaborated on editing and revising the manuscript.

## Acknowledgments

This work was supported by grant 61454 from the John Templeton Foundation.

## Footnotes

1 The double helix was also proposed in Shepard (1982a) under another set of assumptions pertaining to the perceptual uniformity of the diatonic scale.

2 https://www.psynet.dev

3 https://dallinger.readthedocs.io/en/latest/

4 https://tonejs.github.io/

5 https://gitlab.com/computational-audition/sing4me

6 https://scikit-learn.org/stable/

7 https://scipy.org/

## Appendix A Supplementary Figures and Tables

**Table A1.** Demographics of the lower and upper experience non-musician subgroups, as well as the musician groups.

| Task | Group | $N$ | $N_m$ | $N_f$ | $N_o$ | $M_{age}$ | $SD_{age}$ | $M_{yme}$ | $SD_{yme}$ | $Range$ |
| --- | --- | --- | --- | --- | --- | --- | --- | --- | --- | --- |
| Similarity (Mel.) | No Exp. | 102 | 66 | 36 | 0 | 39.8 | 11.6 | 0 | 0 | 0 – 0 |
| Similarity (Mel.) | Some Exp. | 92 | 54 | 37 | 1 | 41.0 | 12.8 | 7.3 | 8.9 | 0.5 – 53 |
| Similarity (Mel.) | Musicians | 41 | 19 | 15 | 7 | 30.0 | 8.3 | 19.9 | 7.9 | 11 – 41 |
| Singing | No Exp. | 50 | 30 | 20 | 0 | 42.5 | 11.0 | 0 | 0 | 0 – 0 |
| Singing | Some Exp. | 49 | 30 | 19 | 0 | 43.8 | 11.3 | 7.6 | 9.8 | 0.5 – 55 |
| Singing | Musicians | 42 | 15 | 26 | 1 | 31.9 | 10.3 | 21.2 | 9.4 | 12 – 46 |
| Similarity (Ton.) | No Exp. | 94 | 57 | 37 | 0 | 40.9 | 10.4 | 0 | 0 | 0 – 0 |
| Similarity (Ton.) | Some Exp. | 92 | 52 | 39 | 1 | 39.5 | 10.1 | 5.7 | 6.0 | 1 – 34 |
| Similarity (Ton.) | Musicians | 30 | 14 | 10 | 6 | 32.4 | 10.6 | 22.2 | 9.6 | 12 – 46 |

Note: The measures are: $N$ total number of participants. $N_m$ , $N_f$ , $N_o$ number of participants that self-identified as male, female, and other, respectively. $M_{age}$ , $SD_{age}$ mean and standard deviation of age. $M_{yme}$ , $SD_{yme}$ , $Range$ mean, standard deviation and range of reported years of musical experience. ‘Similarity (Mel.)’ and ‘Similarity (Ton.)’ correspond to similarity over melodies and isolated tones, respectively.

**Figure A1.**
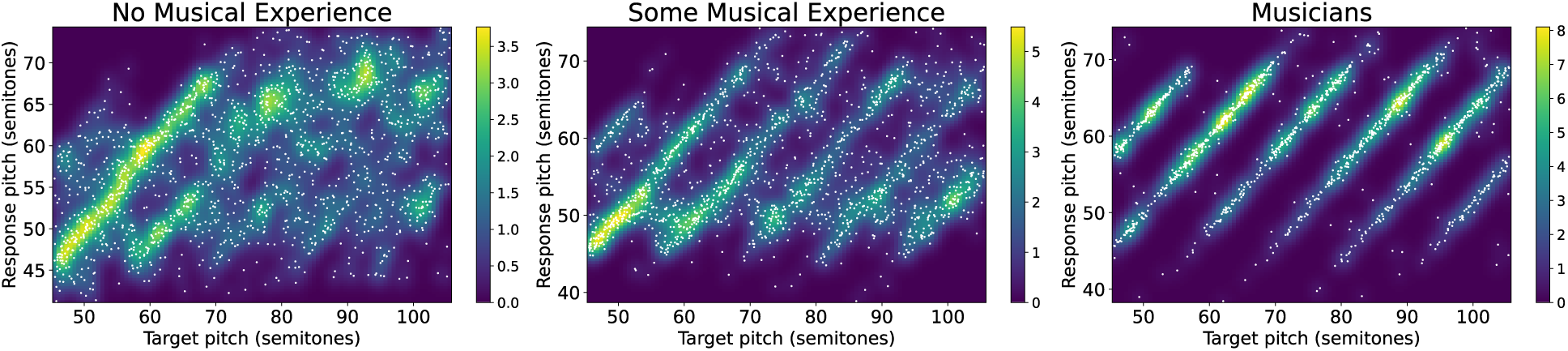
Pitch imitation through singing. Target vs. response pitch distributions for musicians and non-musicians (using a Guassian kernel with σ = 1.1 semitones determined through 5-fold cross validation; density normalized relative to a uniform distribution).

**Figure A2.**
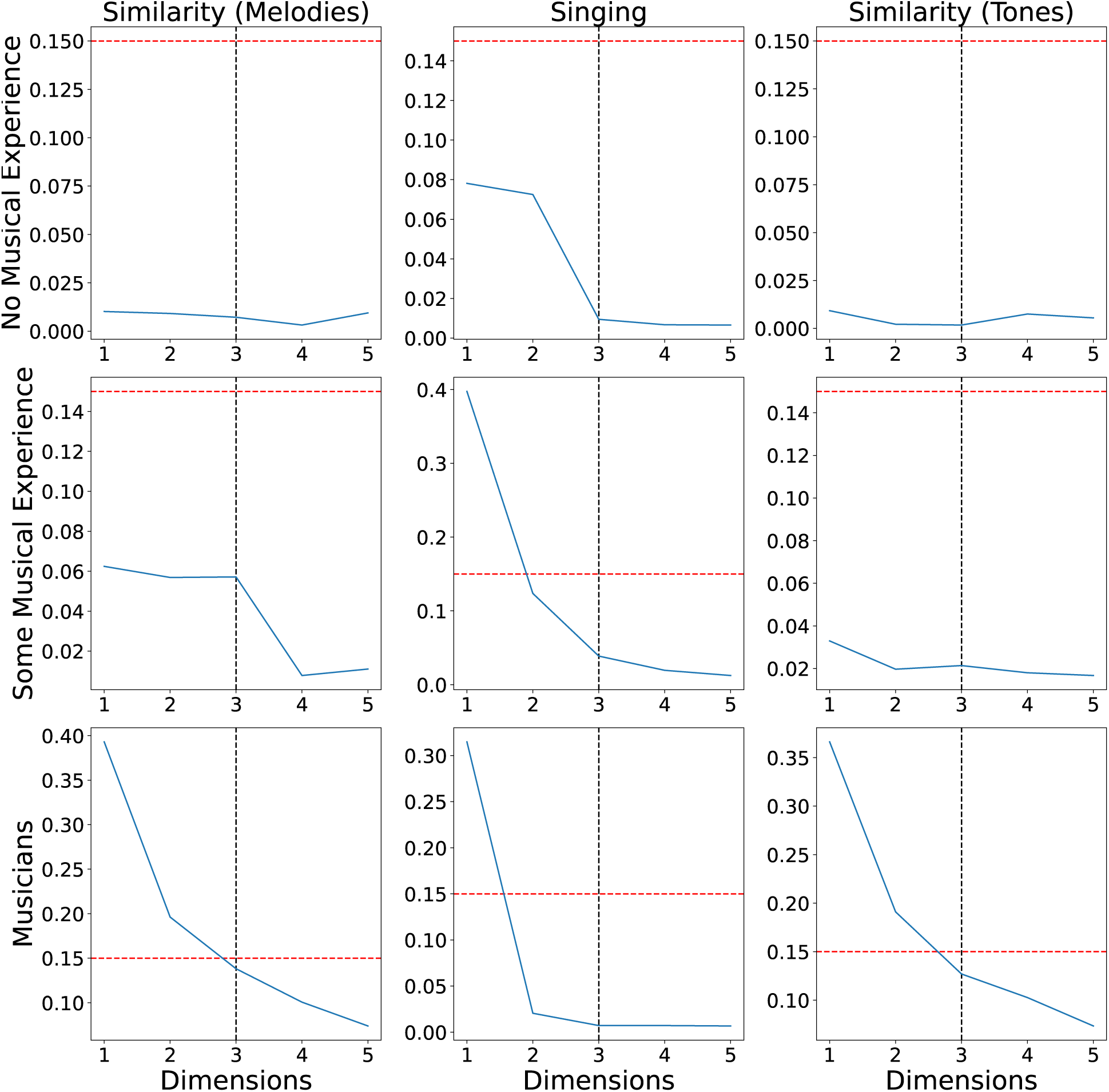
Normalized stress curves as a function of embedding dimensions as applied to the different similarity matrices in Supplementary Figure A4. Dashed red line indicates the 0.15 threshold above which MDS fit is deemed poor (Kruskal, 1964). Based on this analysis, we selected three-dimensional MDS for presenting the results in the paper.

**Figure A3.**
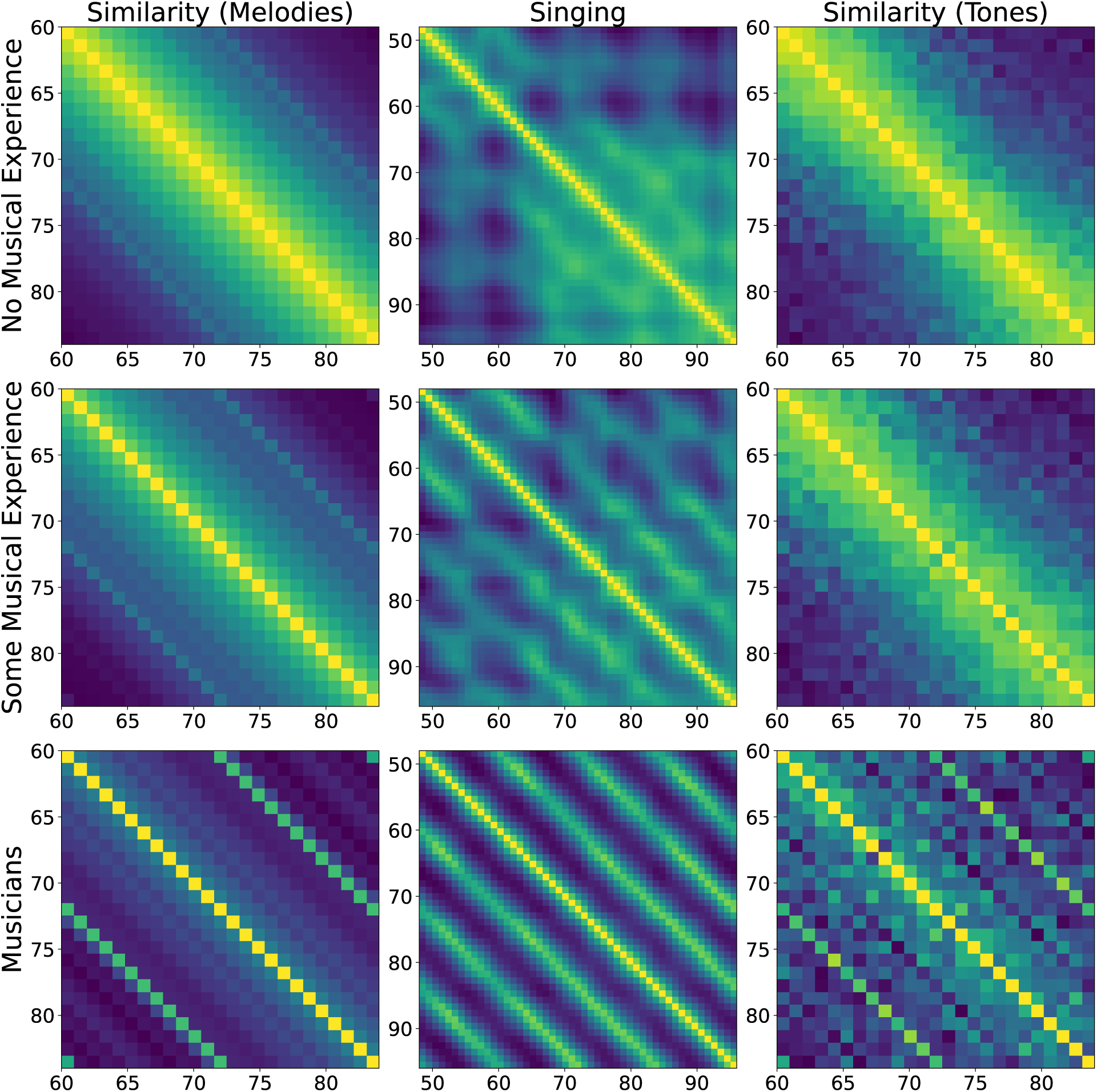
Raw similarity matrices for the different behavioral paradigms considered in the paper.

**Figure A4.**
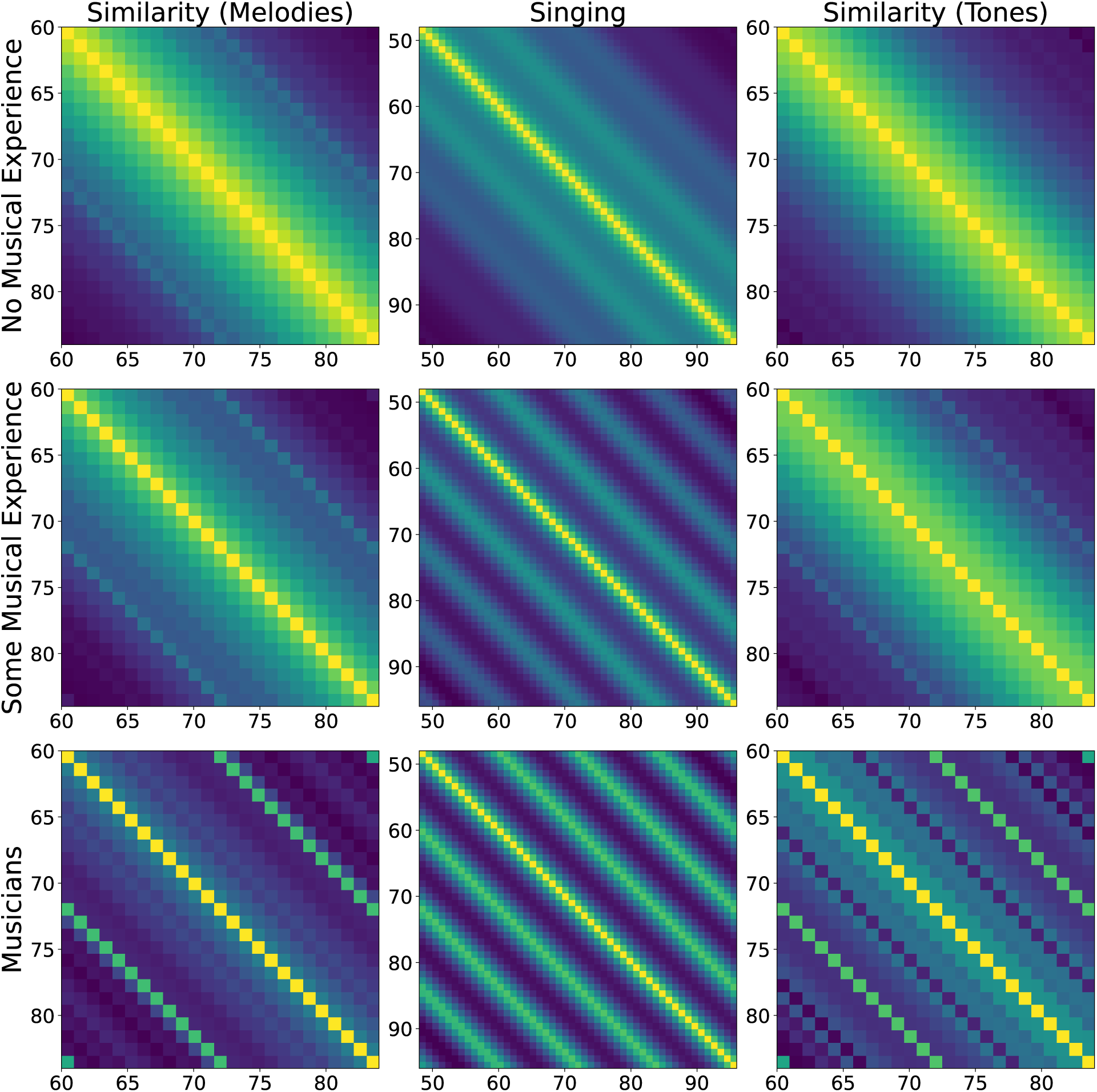
Smoothed similarity matrices by averaging the raw matrices in Figure A3 over their diagonals for the different behavioral paradigms considered in the paper.

**Figure A5.**
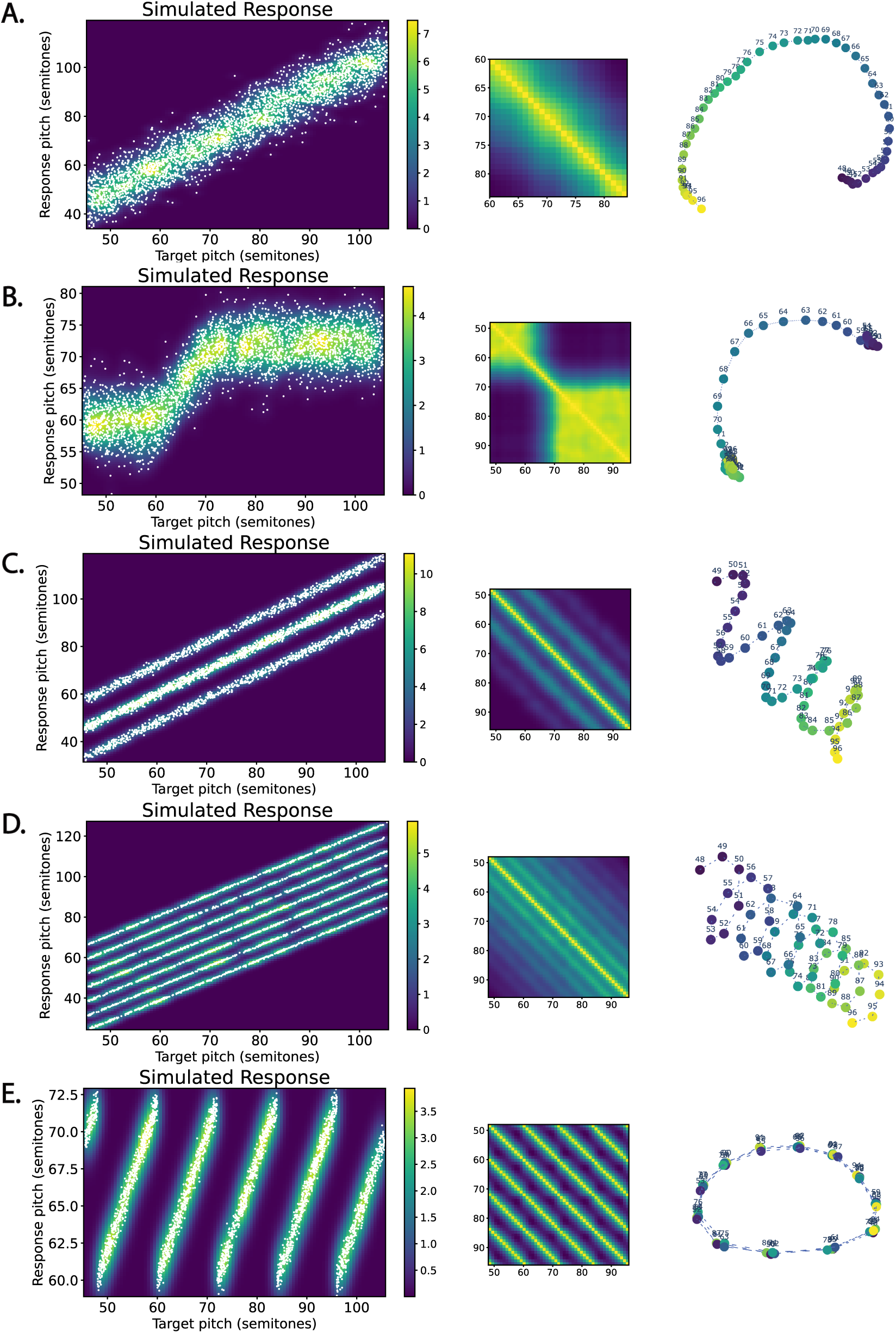
Simulated singing responses and their corresponding similarity matrices and MDS solutions. **A.** Noisy linear response. **B.** Noisy linear response with register saturation. **C.** Singing an octave above or below a tone. **D.** Imitating along the circle of fifths. **E.** Chroma matching.

**Figure A6.**
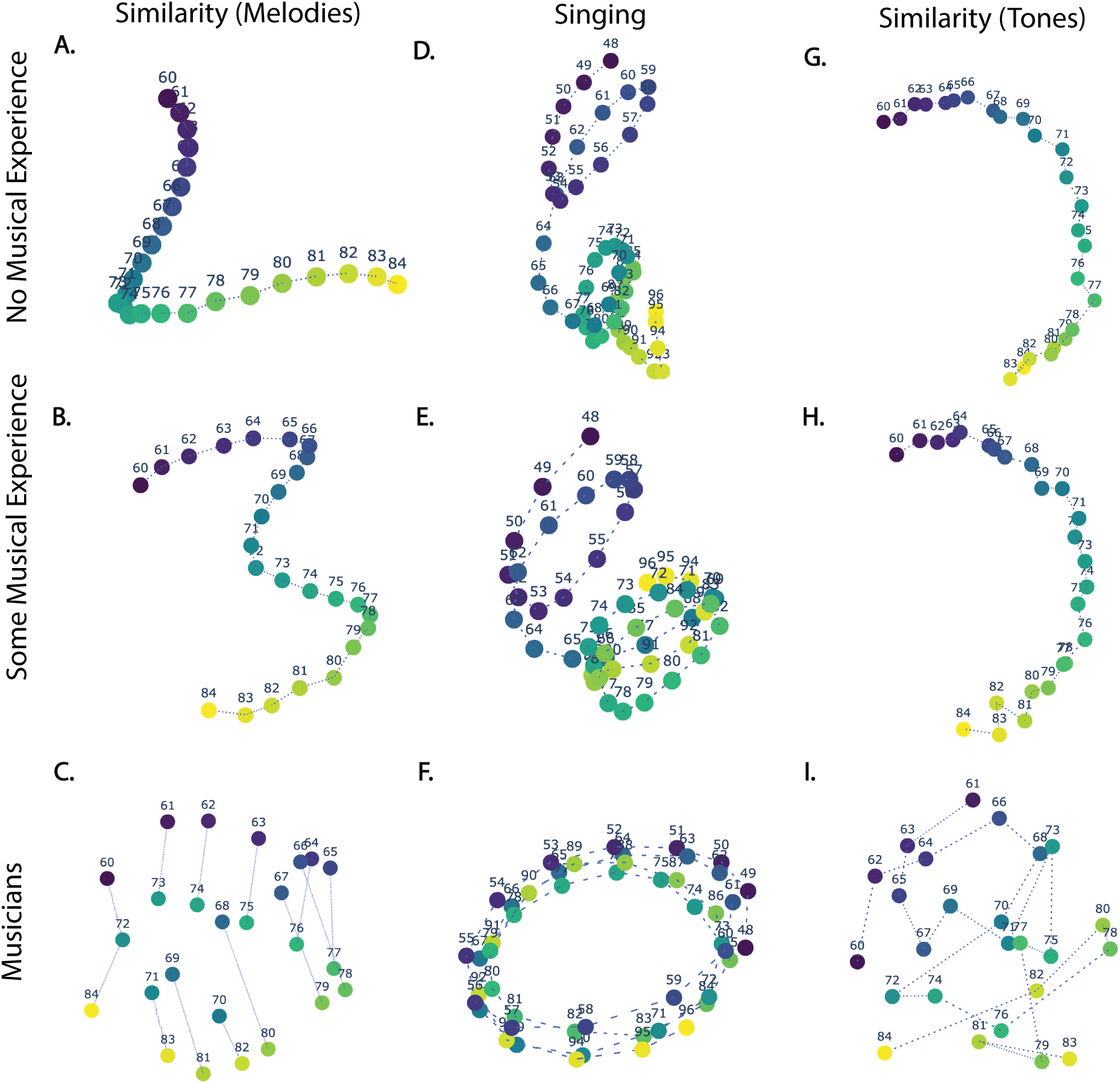
Three-dimensional multidimensional scaling solutions for the raw unprocessed behavioral similarity matrices in Supplementary Figure A3. Left to right: similarity judgments over melodies that differ by a transposition, free imitation of two-note melodies via singing, and similarity judgments over pairs of isolated tones.

**Figure A7.**
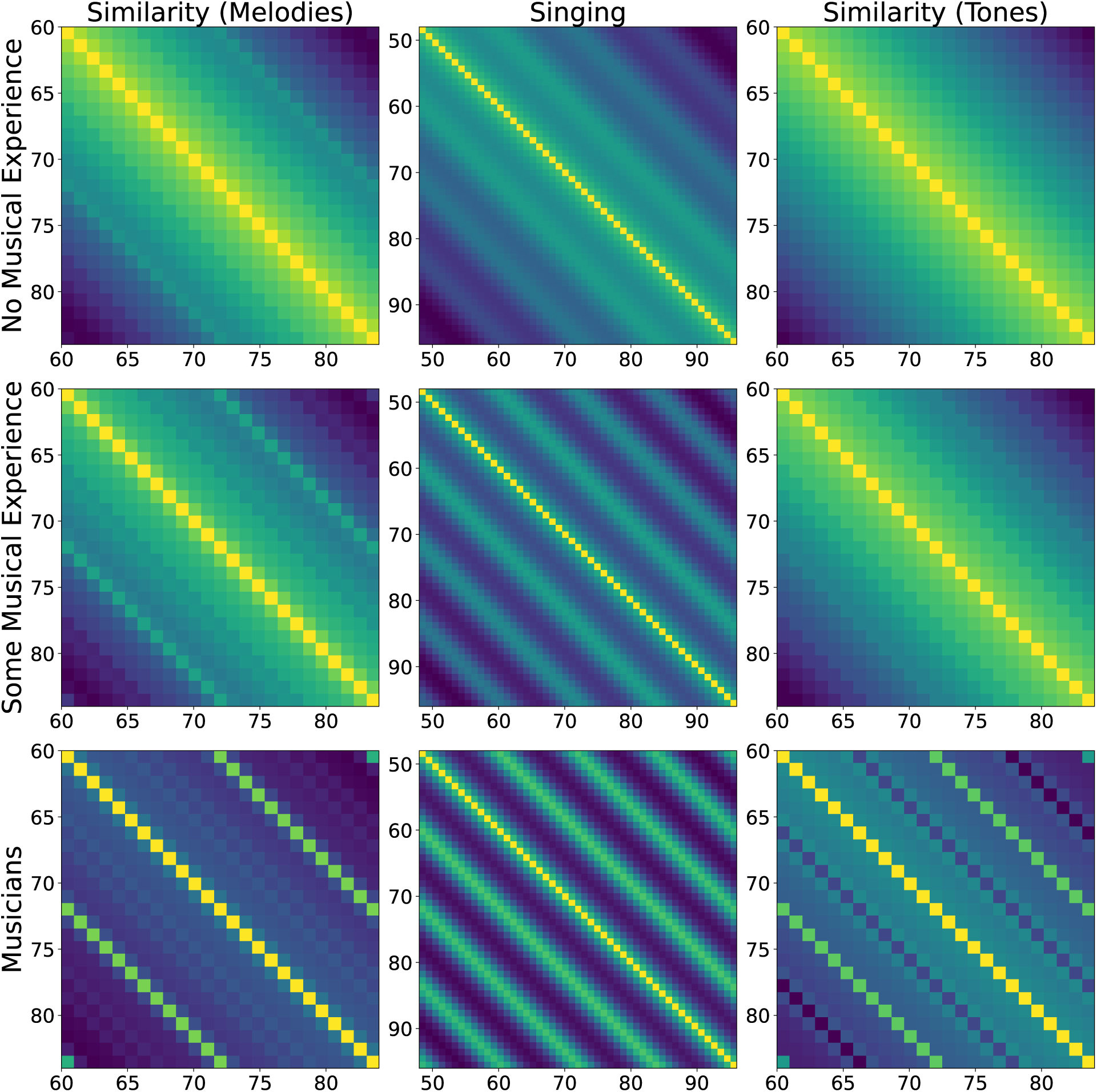
Fitted model similarity matrices for the different behavioral paradigms considered.

**Table A2.**
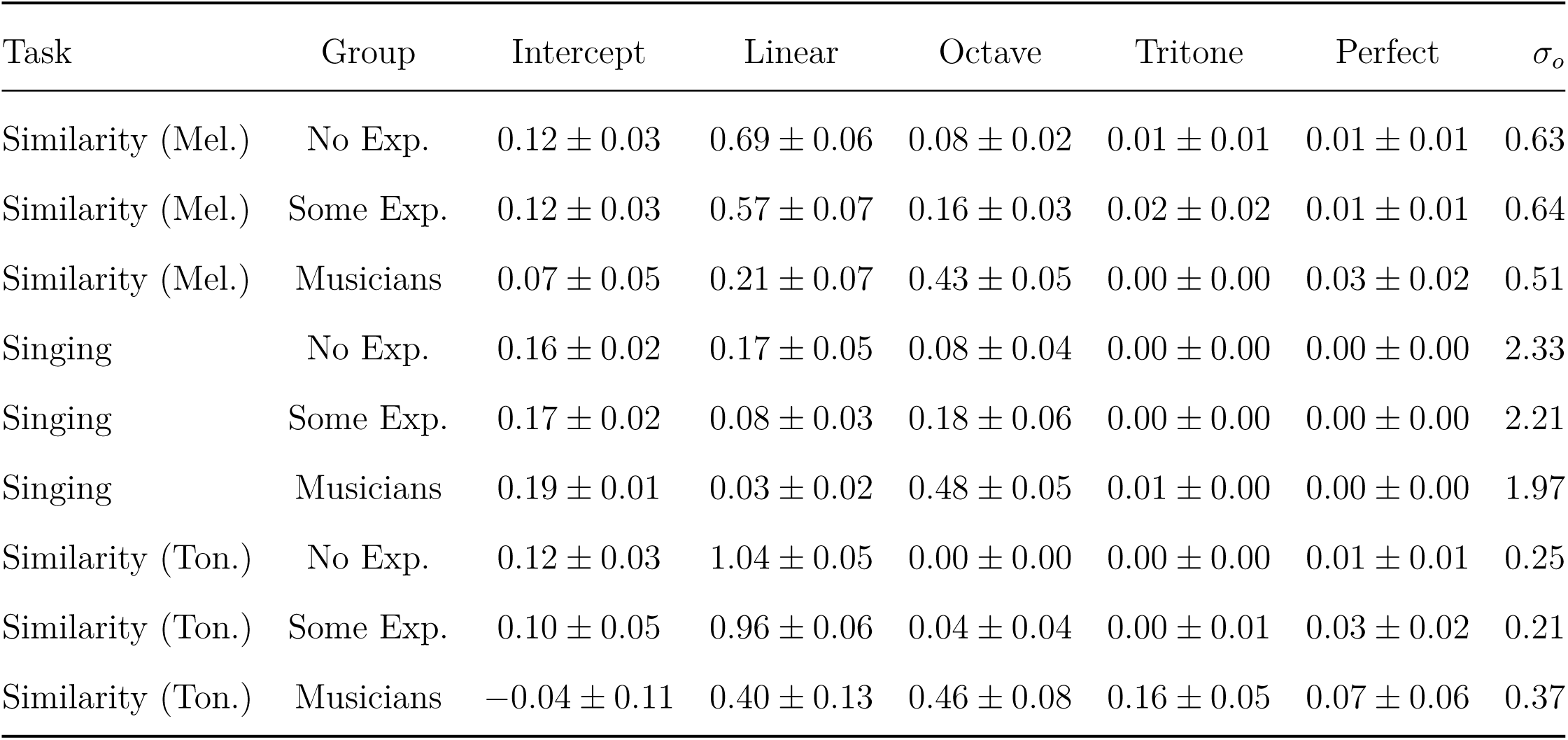
Full list of model parameter values and their 95% confidence intervals.

**Table A3.**
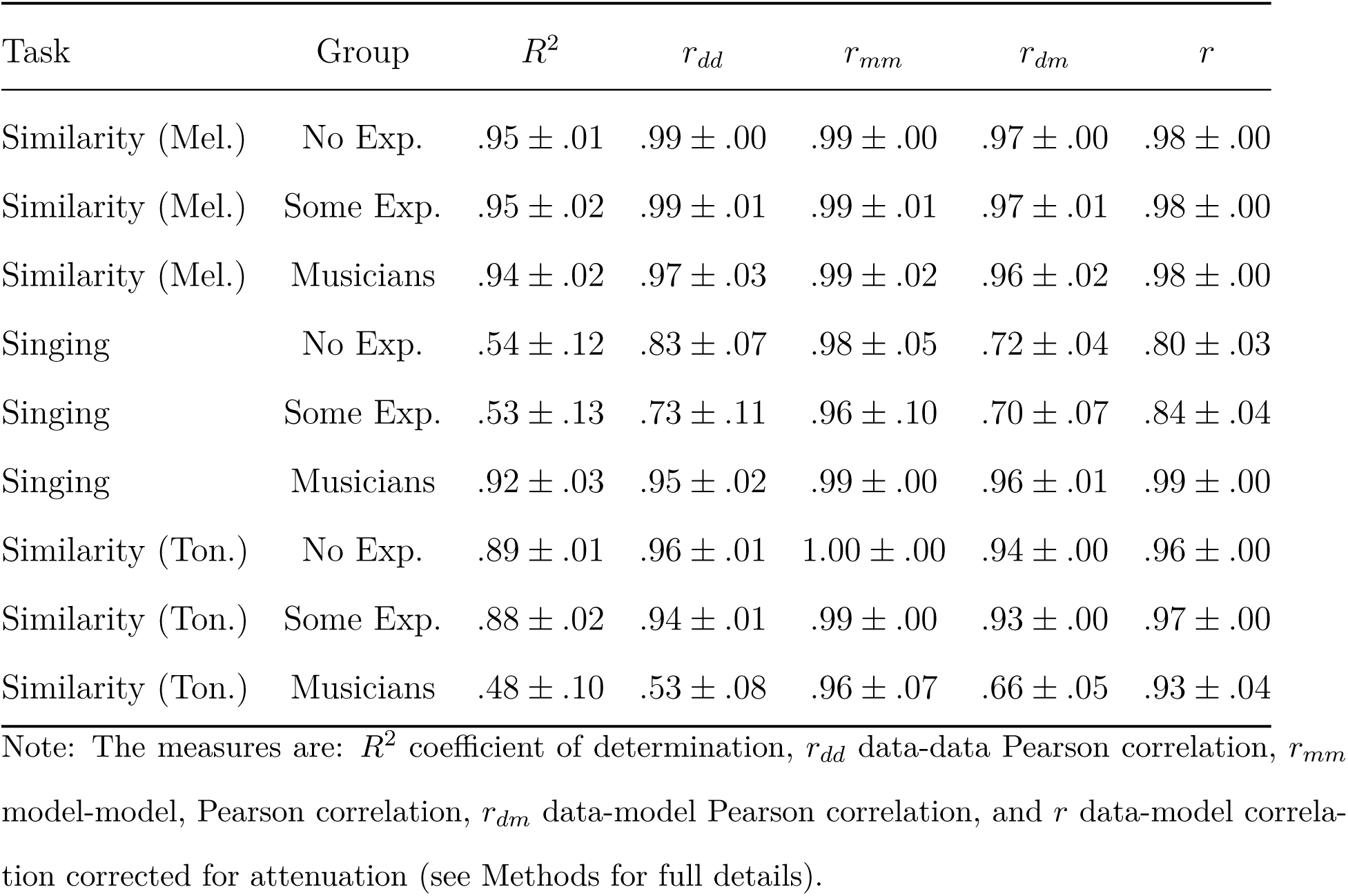
Full list of evaluation metrics and their 95% confidence intervals using split-half bootstrap over participants with 1,000 repetitions.

| Task | Group | $R^2$ | $r_{dd}$ | $r_{mm}$ | $r_{dm}$ | $r$ |
| --- | --- | --- | --- | --- | --- | --- |
| Similarity (Mel.) | No Exp. | .95 ± .01 | .99 ± .00 | .99 ± .00 | .97 ± .00 | .98 ± .00 |
| Similarity (Mel.) | Some Exp. | .95 ± .02 | .99 ± .01 | .99 ± .01 | .97 ± .01 | .98 ± .00 |
| Similarity (Mel.) | Musicians | .94 ± .02 | .97 ± .03 | .99 ± .02 | .96 ± .02 | .98 ± .00 |
| Singing | No Exp. | .54 ± .12 | .83 ± .07 | .98 ± .05 | .72 ± .04 | .80 ± .03 |
| Singing | Some Exp. | .53 ± .13 | .73 ± .11 | .96 ± .10 | .70 ± .07 | .84 ± .04 |
| Singing | Musicians | .92 ± .03 | .95 ± .02 | .99 ± .00 | .96 ± .01 | .99 ± .00 |
| Similarity (Ton.) | No Exp. | .89 ± .01 | .96 ± .01 | 1.00 ± .00 | .94 ± .00 | .96 ± .00 |
| Similarity (Ton.) | Some Exp. | .88 ± .02 | .94 ± .01 | .99 ± .00 | .93 ± .00 | .97 ± .00 |
| Similarity (Ton.) | Musicians | .48 ± .10 | .53 ± .08 | .96 ± .07 | .66 ± .05 | .93 ± .04 |

Note: The measures are: $R^2$ coefficient of determination, $r_{dd}$ data-data Pearson correlation, $r_{mm}$ model-model, Pearson correlation, $r_{dm}$ data-model Pearson correlation, and $r$ data-model correlation corrected for attenuation (see Methods for full details).

**Figure A8.**
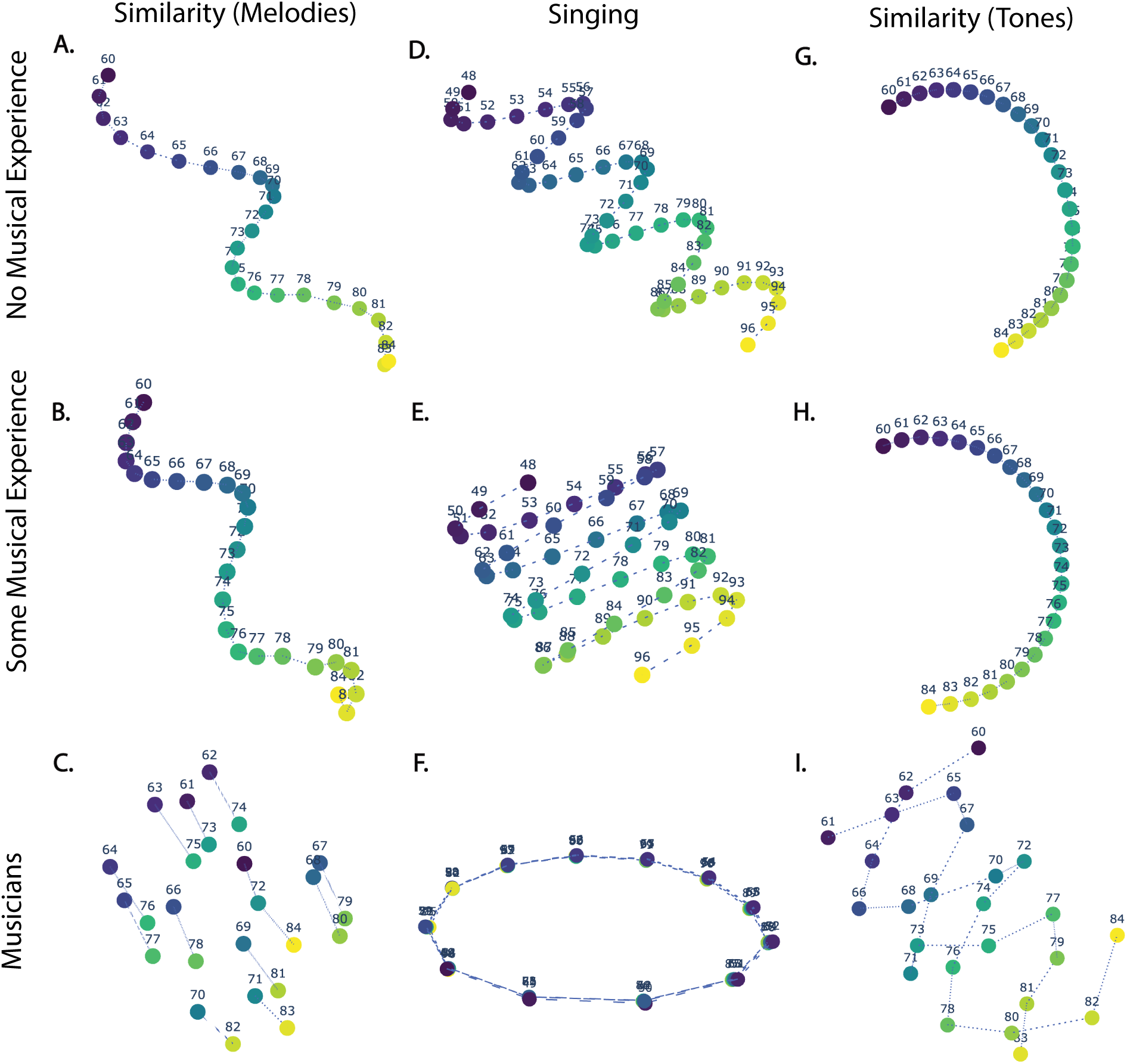
Three-dimensional multidimensional scaling solutions for the fitted model similarity matrices. Left to right: similarity judgments over melodies that differ by a transposition, free imitation of two-note melodies via singing, and similarity judgments over pairs of isolated tones.

**Figure A9.**
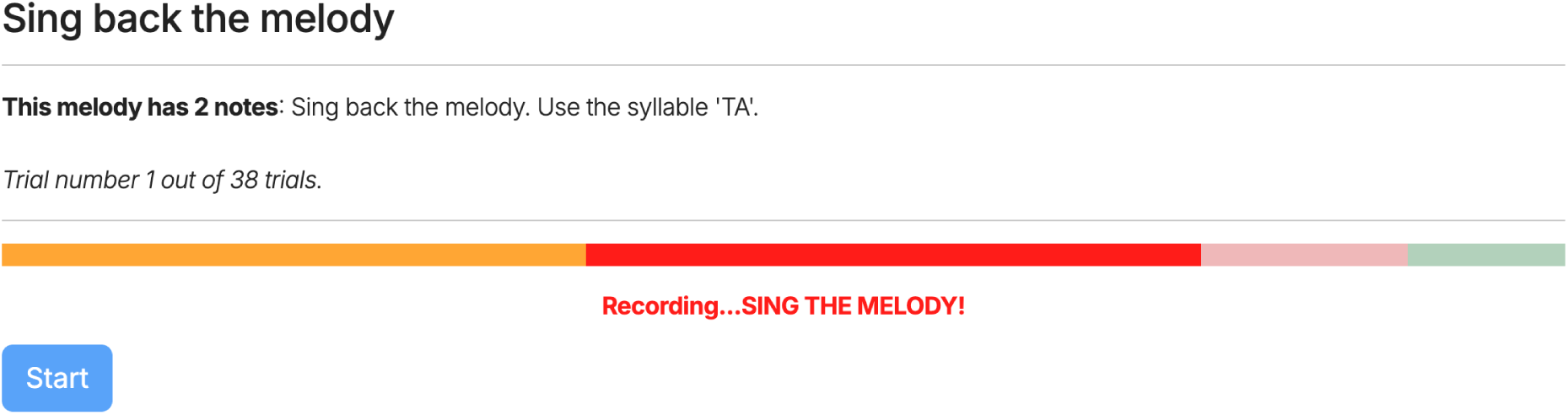
Recording interface for the online singing paradigm.

**Figure A10.**
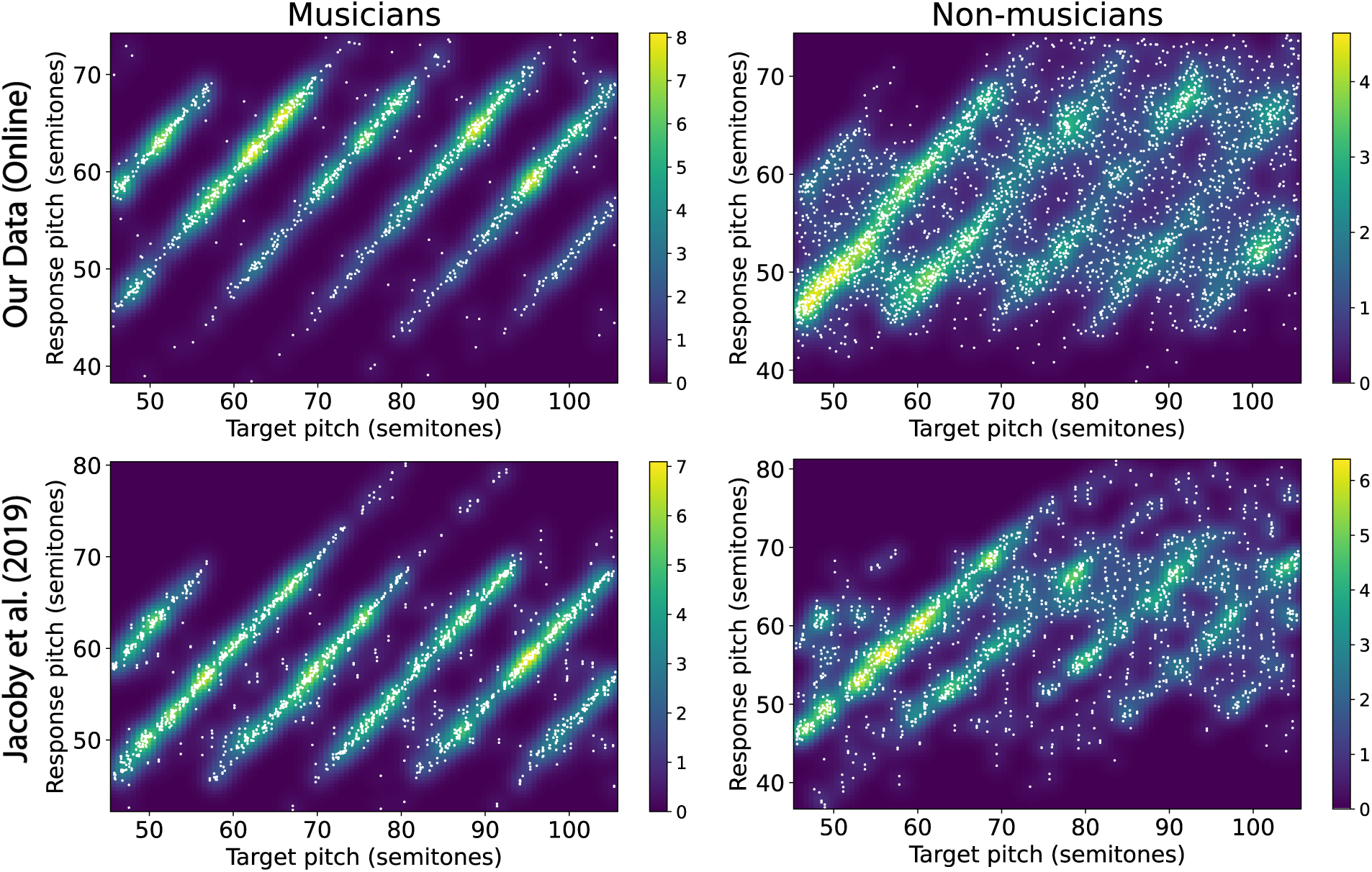
Comparing our online response pitch distributions to the in-lab response distributions of Jacoby et al. (2019) (using a Guassian kernel with σ = 1.1; density normalized relative to a uniform distribution).

**Figure A11.**
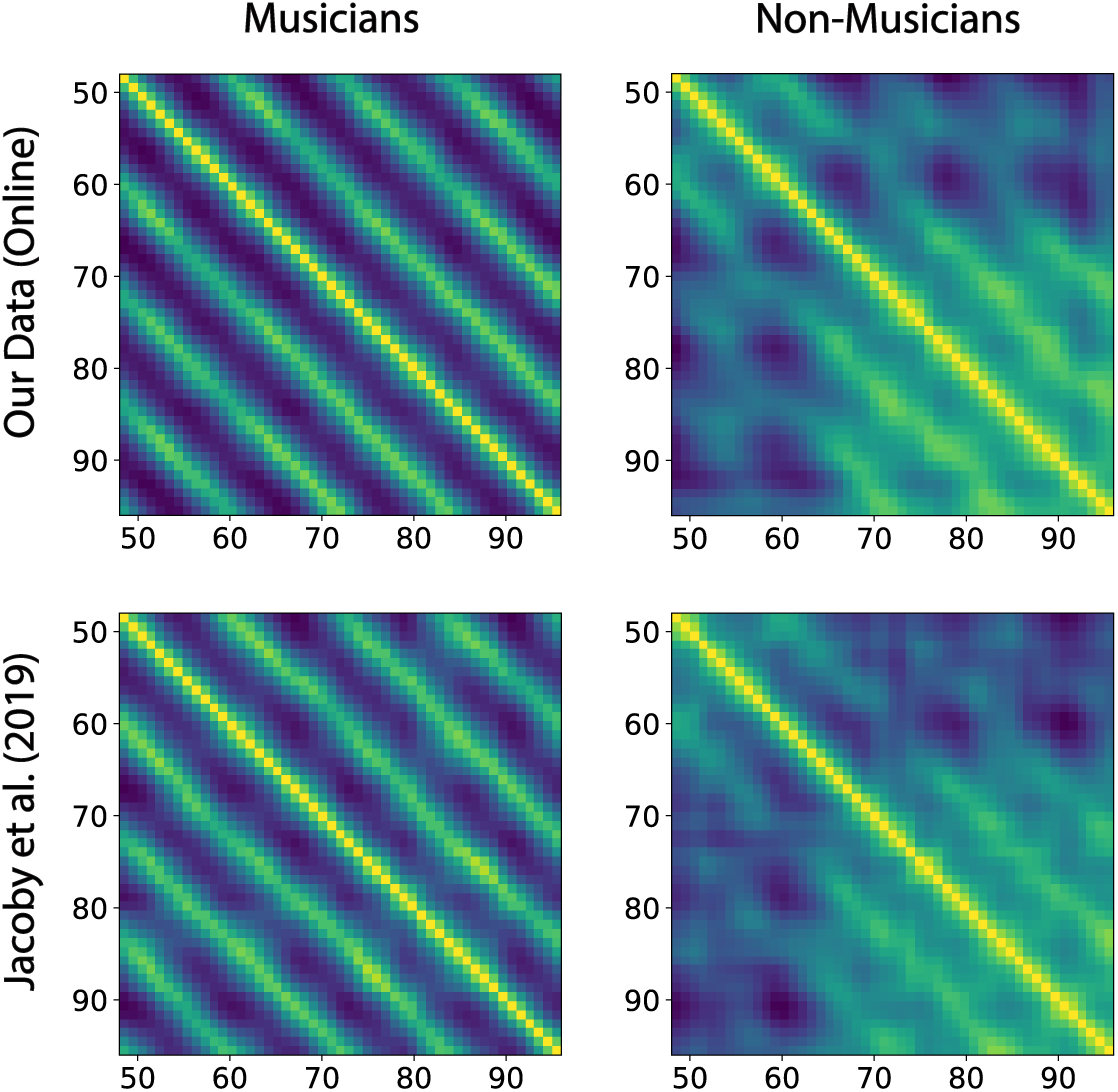
Comparison between resulting similarity matrices based on our online data and the in-lab data of Jacoby et al. (2019) shown in Supplementary Figure A10. The results show high consistency of the similarity matrices between our online data and the in-lab data (r =.93, p < 0.001 for non-musicians, and r =.95, p < 0.001 for musicians)

**Figure A12.**
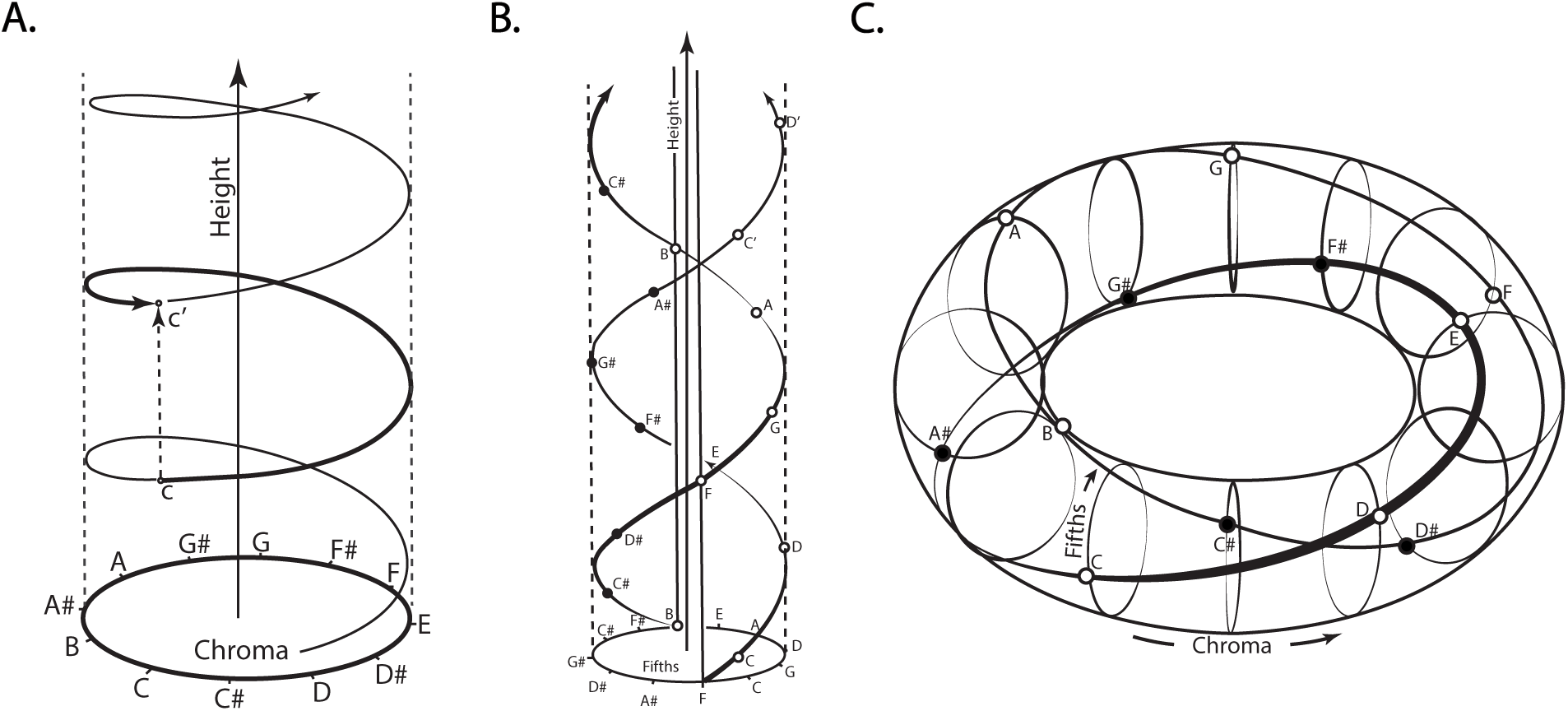
Different geometrical models for the structure of musical pitch as proposed by Roger Shepard. Illustrations are redrawn based on the original designs shown in Shepard (1982a). **A.** A simple helix. **B.** A double helix wound around a line. **C.** A double helix wound around a torus. Musical note values as well as the chroma circle and circle of fifths are indicated.

## Appendix B Additional Experimental Details

### Pre-screening

#### Similarity Paradigms

To enhance online data quality, in the similarity tasks participants were required to complete a headphone prescreening test (Woods et al., 2017) to ensure that they were wearing headphones. In each trial of this test, participants heard a series of three tones and were asked to judge which of these was the quietest. The tones were designed to induce a phase cancellation effect when played on speakers which would lead participants without headphones to answer incorrectly. Participants who failed the headphone test were not allowed to proceed to the main experiment but were nevertheless fully compensated for the time taken to perform the pre-screening test.

#### Singing Paradigm

As with the similarity tasks, participants in the singing paradigm first completed a headphone prescreening test. This was then followed by an additional recording test which was adapted from Anglada-Tort et al. (2023) and proceeded as follows. First, participants performed a microphone check to ensure that their audio was detectable. Specifically, a pop up window was triggered asking participants to allow audio recording in the browser. Participants could not proceed without enabling their microphone. Once allowed, participants were instructed to freely sing into their microphone and check that an audio meter (provided on the page) indicated an appropriate sound level (volume). Next, participants were asked to sing two notes of their choice to the syllable ‘TA’ which were then played back to the participant so that they can double check that their microphone is appropriately set up. At this point, participants completed two practice trials in which they were instructed to imitate two two-note melodies (‘In this task, we want to measure how well you can imitate a melody. In the next few trials you can practice performing the task, before we measure your performance in the main experiment. You will hear a melody with 2 notes, and after it ends, you’ll be asked to sing these two notes back as accurately as possible. Note: Use the syllable TA to sing each note and leave a silent gap between notes. We will provide feedback after each trial’). After each trial participants were informed of the number of notes that were detected (see Singing Transcription Technology below). Importantly, this step did not provide feedback on the singing pitch to avoid biasing the participants. Finally, participants proceeded to a test phase where they had to complete 10 additional singing imitation trials without feedback (‘We will now test your singing performance in a total of 10 trials. Like before, your goal is to listen to each melody, and after it ends, sing these two notes back as accurately as possible. Note: Use the syllable TA to sing each note and leave a silent gap between notes’). Participants recruited through AMT were allowed to proceed only if they were able to satisfy the following three criteria on at least 5 of the 10 trials: (i) They sung exactly two notes, (ii) the absolute interval error between the target and sung melodies was no larger than 3 semitones, and (iii) the direction of the target and sung melodies was the same (i.e. ascending vs. descending). These criteria were adapted from Anglada-Tort et al. (2023) and intended to ensure that AMT participants were making an honest effort to imitate the melodies. Musicians also performed this pre-screener, however, we set their passing threshold to 0 (meaning that everyone passed) as their recruitment was more controlled. Nonetheless, in a post-hoc analysis we confirmed that only one musician marginally failed the test (by achieving a score of 4) whom we excluded. Finally, we note that the two-note melodies were taken from the same set as that devised by Anglada-Tort et al. (2023), and comprised the melodic intervals-2.6,-1.3, 0.0, 1.3, and 2.6, played at two registers (i.e. bass tones), 49 semitones (low) and 61 semitones (high), with uniform roving width of 2.5 semitones. The fractional intervals and roving were applied so that no specific sense of tonality is induced (Jacoby et al., 2019).

#### Performance Incentives

To motivate online participants to provide honest responses for the otherwise subjective tasks, participants in the similarity paradigms were informed that they could receive a performance bonus depending on the quality of their responses. Specifically, they received the following instructions: “The quality of your responses will be automatically monitored, and you will receive a bonus at the end of the experiment in proportion to your quality score. The best way to achieve a high score is to concentrate and give each trial your best attempt”. While the tasks are subjective in nature, we used self-consistency as a measure of performance quality. This was done by repeating 5 random trials at the end of the experiment and then computing Spearman correlation between the original and repeated answers. The resulting score *s* was then used to compute a small bonus of up to 10 cents using the formula min(max(0, 0.1*s*), 0.1).

#### Singing Transcription Technology

We relied on the singing transcription technology^5^ of Anglada-Tort et al. (2023) for extracting the fundamental frequencies (*f*_0_) of the participants’ sung notes. This technology has been extensively tested and was shown to provide reliable *f*_0_ estimates in online settings (Anglada-Tort et al., 2023). The automated process comprises four steps: i) Cleaning the recorded audio from artifacts. This included removing the first 100 ms of a recording, applying a 150 ms linear fade-in, and applying a band-pass filter with cut-off frequencies 80-6000 Hz to remove any non-singing spectral components. ii) Applying a pitch estimation method based on auto-correlation (Boersma et al., 1993) for extracting *f*_0_ from sung segments, implemented using parselmouth (Jadoul et al., 2018). We used the recommended parameters (Anglada-Tort et al., 2023) of 65 Hz (36 semitones, C2) for pitch floor and 622 Hz for pitch ceiling (75 semitones, D#5). iii) Segmenting the continuous *f*_0_ signal into individual isolated tones. This was done by extracting the envelope of the audio signal (by computing the maximum of the squared amplitude within non-overlapping 40 ms windows and then taking the square root) and applying a peak detection algorithm (Duarte & Watanabe, 2015) to find areas of vocal activity that are separated by silences (defined as envelope values that are less than-30 dB relative to the first peak within a segment and extending for at least 30 ms). iv) Filtering the list of audio segments using a set of heuristics to remove any spurious audio segments. This included removing segments that are shorter than 40 ms, removing segments with large deviations from the median *f*_0_, i.e., segments that deviate from the median by at least 8 semitones for more than 35% of the duration the segment which could happen when the singer’s voice “cracks”, as well as removing any segments with median *f*_0_ that falls outside the 65 - 622 Hz range of the auto-correlation step). Finally, the remaining segments were summarized into single pitch values by computing the median *f*_0_ (full details of the approach can be found in Anglada-Tort et al. (2023) under Method Details).

To ensure that the parameters of the singing transcription technology did not bias or constrain the response distributions, we compared our results to the in-lab data of Jacoby et al. (2019). The data are reproduced in Supplementary Figure A10 alongside ours. We see that the online and in-lab data are highly consistent, both in terms of the response range (95% CI of response pitch values for the in-lab data was [44, 76] semitones for non-musicians, and [45, 72] semitones for musicians, both of which are highly overlapping with our design range of [36, 75] semitones), as well as the resulting similarity matrices shown in Supplementary Figure A11 (*r* =.93, *p <* 0.001 for non-musicians, and *r* =.95, *p <* 0.001 for musicians; see below for details regarding computing the similarity matrices).

#### Singing Response Distributions

To construct similarity matrices based on the response distributions of human singers, we first applied a 2D Gaussian kernel density estimate (KDE) *ρ*(*p_t_, p_r_*) to the target-response pitch pairs (*p_t_, p_r_*) (we used the first tone in each target melody as the second tone is not independent) using the KernelDensity method of the scikit-learn^6^ Python package with a bandwidth parameter of *σ* = 1.1 semitones and a resolution of 500 × 500 bins. We used this bandwidth as it provided a good tradeoff between reliability and resolution as determined by a five-fold cross validation analysis using the GridSearchCV method of scikit-learn. Dissimilarity between two target pitches *p_t_*_1_ and *p_t_*_2_ was then computed by applying the Jensen-Shannon distance jensenshannon from the scipy^7^ package to the KDE marginals *ρ*(*p_r_*|*p_t_*_1_) and *ρ*(*p_r_*|*p_t_*_2_). To ensure that this approach is sufficiently flexible to capture different meaningful structures we applied it to different simulated response distributions that capture various modes of behavior. Specifically, we used a similar target pitch distribution as in the online experiments and simulated different response pitch values including (i) a noisy linear response, (ii) a noisy linear response with register saturation, (iii) a mixture of linear response with some probability of singing an octave above or below the target pitch, (iv) a mixture of linear response with some chance of singing one or more fifths above or below the target pitch, and (v) chroma matching within a given register. We then applied an identical processing procedure as in the behavioral experiments to produce similarity matrices which we then subjected to MDS. The results are shown in Supplementary Figure A5 and we found that the approach is indeed able to capture a variety of qualitatively different structures. Simulation code and 3D visualizations of the simulated solutions are provided in the OSF repository.

